# Dynamics and composition of small heat shock protein condensates and aggregates

**DOI:** 10.1101/2022.12.08.519563

**Authors:** Joep Joosten, Bob van Sluijs, Wilma Vree Egberts, Martin Emmaneel, Pascal W.T.C. Jansen, Michiel Vermeulen, Wilbert Boelens, Kimberly M. Bonger, Evan Spruijt

## Abstract

Small heat shock proteins (sHSPs) are essential ATP-independent chaperones that protect the cellular proteome during stress. These proteins assemble into polydisperse oligomeric structures, the composition of which dramatically affects their chaperone activity. The biomolecular consequences of variations in sHSP ratios, especially inside living cells, remain elusive. Here, we study the consequences of altering the relative expression levels of HspB2 and HspB3. These chaperones are partners in a hetero-oligomeric complex, and genetic mutations that abolish their mutual interaction are associated with myopathic disorders.

HspB2 displays three distinct phenotypes when co-expressed with HspB3 at varying ratios. Expression of HspB2 alone lead to formation of liquid nuclear condensates, while shifting the stoichiometry towards HspB3 resulted in the formation of large solid-like aggregates. Only cells co-expressing HspB2 with a limited amount of HspB3 showed a homogeneous nuclear distribution of HspB2. Strikingly, both condensates and aggregates were reversible, as shifting the HspB2:HspB3 balance in situ resulted in dissolution of these structures.

To uncover the molecular composition of HspB2 condensates and aggregates, we used APEX-mediated proximity labelling. Most proteins interact transiently with the condensates and were neither enriched nor depleted. In contrast, we found that HspB2:HspB3 aggregates sequestered several disordered proteins among which autophagy factors, suggesting that the cell is actively attempting to clear these aggregates. This study presents a striking example of how changes in the relative expression levels of interacting proteins affects their phase behavior. Our approach can be a useful tool to study the role of protein stoichiometry in other biomolecular condensates.

**Graphical abstract:** 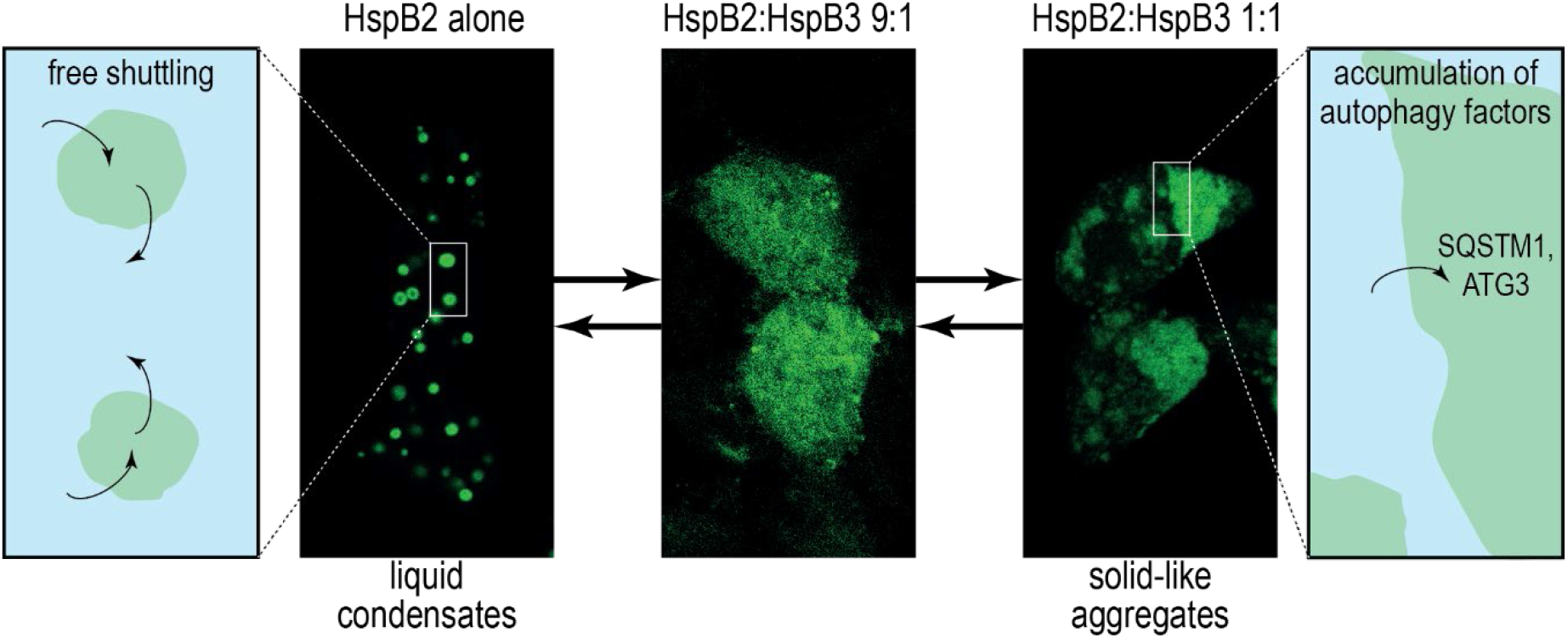

**Highlights:**

- Small heat shock protein hetero-oligomerization affects their chaperone function
- The HspB2:HspB3 expression ratio determines phase separation and aggregation
- HspB2 condensates and HspB2:HspB3 aggregates are fully reversible
- Proximity labelling unveils autophagy factor recruitment to HspB2:HspB3 aggregates
- Stoichiometry-dependant regulation of phase behaviour may be widespread in biology

## INTRODUCTION

Small heat shock proteins (sHSPs) are ATP-independent molecular chaperones that are part of the cellular protein quality control system which defends the integrity of the proteome during cellular stress (1). sHSPs have the ability to bind a large variety of non-native and misfolded proteins, thereby delaying the formation of irreversible protein aggregates (2). After being bound by sHSPs, misfolded proteins are either degraded by the proteasome or autophagosomes, or refolded by ATP-dependent chaperones, such as HSP70 (3–7). As misfolding and aberrant aggregation of proteins are central to the etiology of numerous degenerative diseases (8), it is not surprising that mutations in sHSP genes have been linked to various pathological conditions (9).

The human genome encodes ten sHSPs (HspB1-HspB10), which strongly vary in their tissue expression pattern and levels (Figure S1A-B). Four sHSPs (HspB1, HspB5, HspB6, and HspB8) are ubiquitously expressed at relatively high levels (10), while others are predominantly expressed in specific tissues (Figure S1A-D). For instance, HspB2 (also known as MKBP) and HspB3 (also known as HspL27) are upregulated during myoblast differentiation (11) and highly expressed in heart and skeletal muscle (Figure S1C) (11–13). All sHSPs contain a highly conserved α-crystallin domain (ACD) at their core, which is responsible for the formation of dimers that may ultimately assemble into polydisperse oligomeric complexes up to 1000 kDa in size (14). The ACD is flanked by less conserved, flexible N- and C-terminal regions, which are important for the stabilization of these oligomeric complexes (15,16), and may also mediate phase separation (17).

A recent study has shown that in differentiating myoblasts, HspB2 forms both nuclear foci that do not contain HspB3, as well as cytoplasmic foci that colocalize with HspB3 (18). These findings indicate that, in vivo, HspB2 and HspB3 can form multimeric assemblies of diverse compositions. The dynamics, molecular composition and functional significance of these different structures is currently unknown. By contrast, in vitro, it has been shown that HspB2 and HspB3 form a hetero-tetrameric complex with a well-defined 3:1 ratio that is stable in solution (19,20). While it remains unclear if these chaperones interact in the same way in living cells, it is likely that the balance of expression between the two proteins is important for their subcellular distribution and chaperone activity (21). Thus far, the effects of variation in relative HspB2:HspB3 expression levels on complex formation and distribution in livings cells remains enigmatic (18). Identification of potential sequestered proteins may help uncover the role of foci formed by small heat shock proteins, thereby further elucidating their function, which may shed light on their role in the development of degenerative diseases.

Here, we show that HspB2 forms nuclear foci through liquid-liquid phase separation in the absence of HspB3. Co-expression of a limited amount of HspB3 resulted in dissolution of HspB2 condensates and homogeneous distribution of HspB2 in the nucleus, while further shifting the balance towards HspB3 resulted in the formation of nuclear and cytoplasmic aggregates. The cytoplasmic aggregates in particular are very large and dramatically impact nuclear morphology. These findings underscore the importance of balanced expression of the two chaperones. To elucidate the composition of HspB2 condensates and HspB2:HspB3 aggregates, we used proximity labeling mediated by the engineered peroxidase APEX (22). We found that HspB2:HspB3 aggregates were highly enriched for core autophagy factors, such as SQSTM1/p62, suggesting that these aggregates are targeted by the autophagosome. In contrast, very few proteins were strongly enriched or depleted in nuclear HspB2 condensates, suggesting that most proteins are able to transiently shuttle in and out of the condensates, and that these structures are well tolerated by cells.

## RESULTS

### HspB2 forms condensates through liquid-liquid phase separation

As a member of the sHSP family (16), HspB2 contains the characteristic α-crystallin domain (ACD) domain, flanked by less structurally defined N- and C-terminal tails (Figure 1A). Moreover, the C-terminal tail is highly enriched in negatively charged amino acids, including a repeat of five (E) glutamic acid residues (Figure 1A). In recent years, it has become evident that disordered regions in proteins can serve important biological functions (23,24) and may mediate the formation of condensates through liquid-liquid phase separation (25). Indeed, it has been shown that HspB2 phase separates into nuclear condensates in differentiating myoblasts (18).

**Figure 1.**
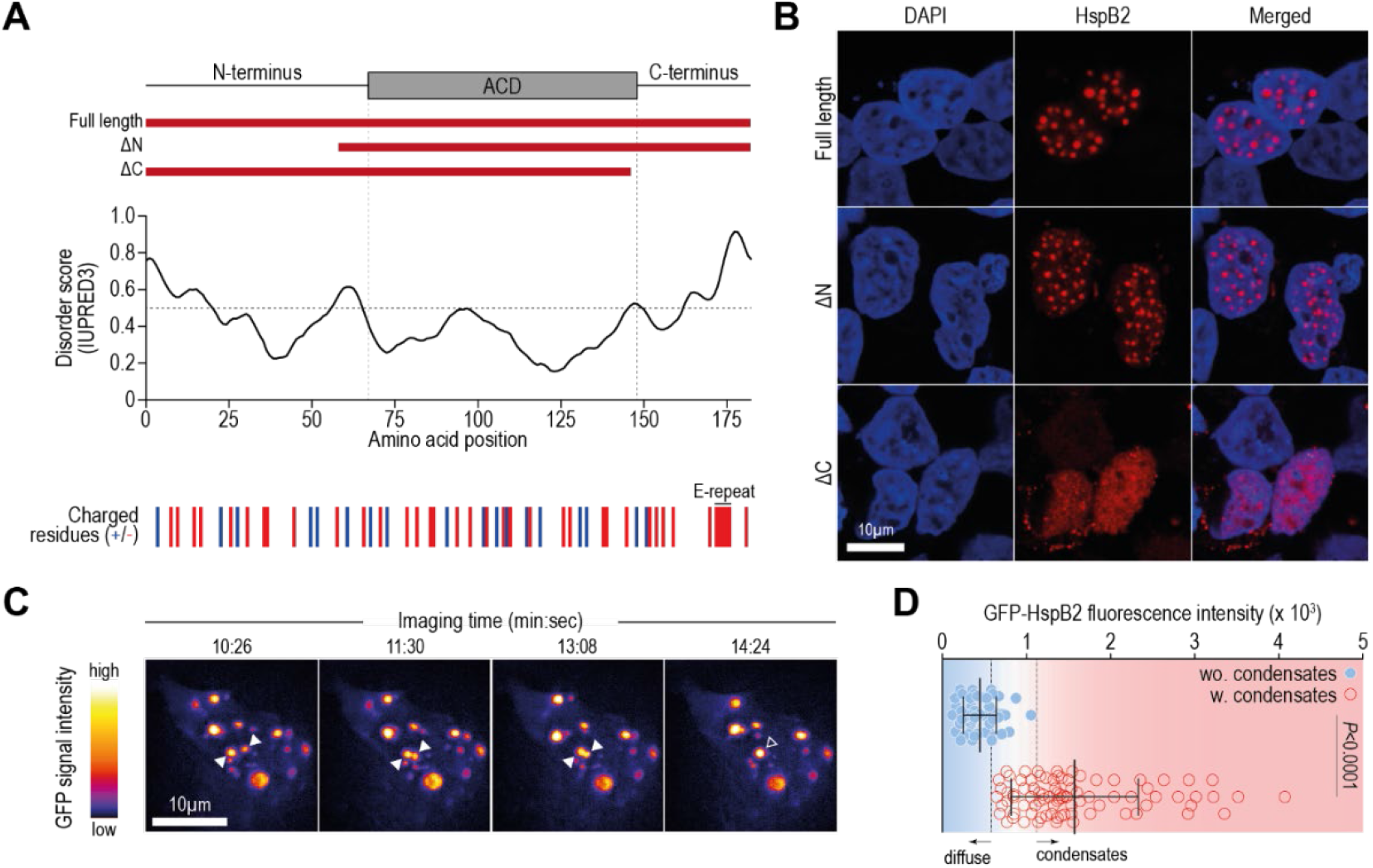
HspB2 condensation into nuclear liquid droplets is regulated by its disordered C-terminus. **(A)** Schematic representation of the HspB2 protein, with the α-crystallin domain (ACD) indicated in grey. Indicated below are the truncation mutants ΔN and ΔC used in (B). The disorder score was determined using IUPRED3. On the bottom, the position of positively (blue; lysine, arginine, and histidine) and negatively (red; aspartic acids and glutamic acid) charged amino acid residues is indicated. **(B)** Confocal microscopy images of HEK293 cells expression transgenic constructs depicted in (A), stained using an antibody against HspB2. The scalebar represents 10 µm. **(C)** Screenshots taken from a video of a HEK293 cell expressing (GFP)-HspB2. Imaging started ~16 hours after transfection and continued for ~15 minutes with an image being taken every 2 seconds. Droplets that will undergo fusion are indicated by closed arrowheads, the fused droplet is indicated by an open arrowhead. The scalebar represents 10 µm. **(D)** Average GFP-HspB2 fluorescence intensity distribution of cells transfected with GFP-HspB2 and untagged HspB2 at a 1:8 ratio. The diffuse (non-condensate) GFP-HspB2 signal was quantified in cells in which condensates were formed (red) and in cells lacking observable condensates (blue). Line and whiskers indicate the mean and standard deviation. A Kolmogorov-Smirnov non-parametric test was performed to determine statistical significance. See also Figure S2 and Videos S1 and S2.

Here, we opted to use HEK293 cells to study the effects of altering the relative HspB2:HspB3 expression levels, as these cells do not express HspB2 and HspB3 endogenously, thereby allowing full control over their relative expression levels. We found that HspB2 forms nuclear foci when overexpressed in HEK293 cells (Figure 1B). To assess the contribution of the disordered N- and C-termini to the formation of these nuclear HspB2 foci, we generated truncation mutants lacking these tails (schematically shown in Figure 1A). While removal of the N-terminus (ΔN) did not affect HspB2 localization, removal of the negatively charged C-terminal tail (ΔC) abolished the formation of nuclear foci, and resulted in a diffuse nuclear distribution of the truncated HspB2 protein (Figure 1B), similar to what was observed previously in differentiating myoblasts (18).

The circularity of the nuclear foci indicates a liquid state, suggesting that liquid-liquid phase separation may underlie their formation. To assess the liquid nature of these foci, we performed live imaging of HEK293 cells expressing GFP-tagged HspB2. We transfected GFP-HspB2 together with untagged HspB2 at a 1:8 ratio to minimize the potential effect of the GFP-tag on the phase behavior of HspB2. At this ratio, the GFP-HspB2 signal recapitulated the nuclear foci previously observed for untagged HspB2 alone (Figure S2A). During live imaging, we predominantly observed dynamically moving nuclear droplets, as well as diffuse cytoplasmic signal and several small, fast-moving cytoplasmic droplets (Video S1). The nuclear GFP-HspB2 droplets fuse upon collision (Figure 1C and Video S2), proving their liquid nature. As condensate formation through liquid-liquid phase separation depends on protein concentration (25), we assessed the average fluorescence intensity of the diffuse (non-condensate) signal in cells that contain HspB2 condensates versus cells lacking condensates. We found that condensates form above a specific concentration threshold (Figure 1D), further supporting the notion that these foci are indeed formed through liquid-liquid phase separation. Together, these data show that HspB2 condensates into nuclear droplets through liquid-liquid phase separation, and that the disordered C-terminus is required for this phase separation.

### Co-expression of HspB3 disrupts the formation of HspB2 condensates

As HspB2 directly interacts with HspB3 (18–20), we assessed the effect of HspB3 co-expression on the subcellular localization and phase behavior of HspB2. We transfected HspB2 with varying amounts of HspB3 into HEK293 cells, again using GFP-tagged HspB2 and untagged HspB2 in a 1:8 ratio to enable live imaging. Similar to our findings using HspB2 antibody staining (Figure 1B), GFP-HspB2 localized to nuclear condensates in the absence of HspB3 (Figure 2A). Co-expression of a small amount of HspB3 (HspB2:HspB3 9:1) resulted in a diffuse distribution of HspB2, mostly in the nucleus. Interestingly however, we observed that further increasing the relative amount of HspB3 (HspB2:HspB3 1:1) resulted in the localization of HspB2 to amorphous aggregates. Both the small HspB2 condensates that form in the absence of HspB3, as well as the nuclear aggregates formed upon co-expression of HspB3 displace chromatin (Figure 2B), potentially affecting gene expression and other nuclear processes. Nonetheless, we did not observe obvious detrimental effects on cell survival or proliferation for any of the HspB2:HspB3 expression ratios used (Figure S2B).

**Figure 2.**
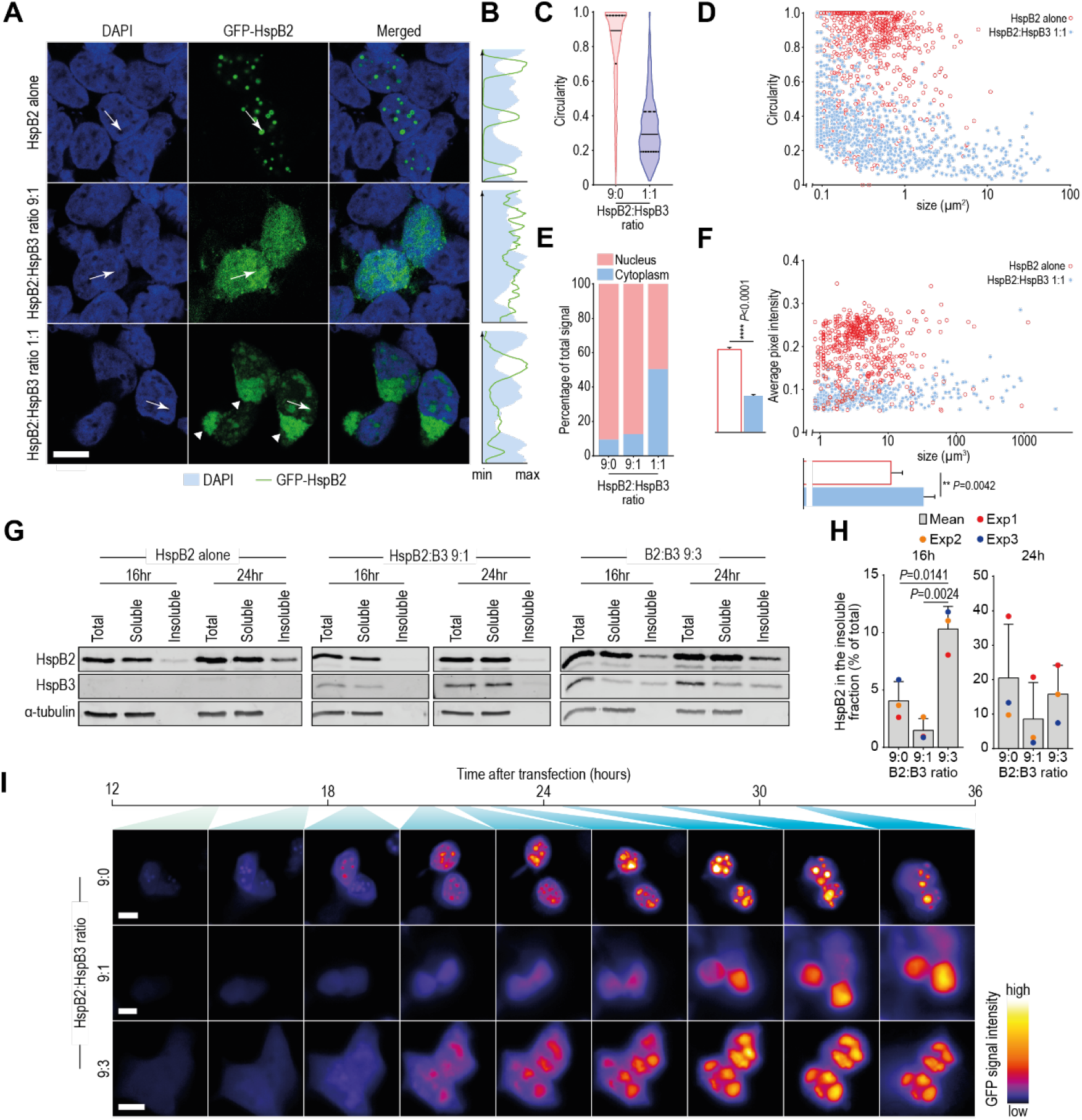
Co-expression of HspB3 disrupts HspB2 phase separation and leads to protein aggregation. **(A)** Confocal images of HEK293 cells transfected with (GFP-)HspB2 and HspB3-expression plasmids at indicated ratios. The scalebar represents 10 µm, arrowheads indicate cytoplasmic aggregates and the arrows designate the direction along which the histograms shown in (B) were determined. **(B)** Histograms depicting GFP-HspB2 and DAPI signal intensities along the lines shown in (A). **(C)** Violin plots showing the circularity of HspB2 foci in cells transfected with HspB2 and HspB3 expression plasmids at indicated ratios. **(D)** Scatter plot depicting the size (x-axis) and circularity (y-axis) of HspB2 foci in cells expressing HspB2 alone, as well as with co-expression of an equimolar amount of HspB3 (HspB2:HspB3 1:1). **(E)** The percentage of pixels located in the nucleus and cytoplasm in cells transfected with HspB2 and HspB3 at indicated ratios. **(F)** Scatter plot depicting the size (x-axis) and average pixel intensities (y-axis) for HspB2-foci in cells expressing only HspB2 or **(A)** HspB2 and HspB3 at a 1:1 ratio. Bars and whiskers represent the mean and 95% confidence interval, non-parametric Kolmogorov-Smirnov tests were used to determine statistical significance. The width of the bars is proportional to the number of HspB2-foci identified in each dataset. **(G)** Western blot analysis depicting the solubility of HspB2 and HspB3 expressed at indicated ratios. Cells were lysed 16 and 24 hours after transfection and fractionated into total, soluble (supernatant) and insoluble (pellet) fractions by centrifugation. **(H)** Quantification of three replicate western blotting experiments. The HspB2 signals were quantified, normalized to the α-tubulin signal in the total lysate sample from the corresponding transfection, and expressed as a percentage of total HspB2. Bars and whiskers are the mean and standard deviation of three biological replicates, colored dots indicate the values from individual experiments. The western blot in (G) is from experiment #3, see also Supplemental Dataset 1 for the raw data underlying this bar graph. Student’s t-tests with correction for multiple testing using the Holm-Šídák method was performed to determine statistical significance. **(I)** Screen shots from videos of HEK293 cells transfected with (GFP)-HspB2 and HspB3 at indicated ratios. In all cases, GFP-HspB2 and untagged HspB2 are expressed at a 1:8 ratio. Imaging started approximately 12 hours after transfection, with an image being taken every 15 minutes for 24 hours. Blue shading corresponds to the position in the timeline shown above. All scalebars represent 10 µm. See also Figure S2, S3 and Videos S3A-C.

Besides nuclear aggregates, additional cytoplasmic structures are formed upon co-expression of HspB2:HspB3 at a 1:1 ratio (indicated with arrowheads in Figure 2A). To verify that these large structures are indeed cytoplasmic, we combined HspB2 imaging with staining of Lamin A and Lamin B1; structural components of the nuclear lamina which line the inner membrane of the nuclear envelope (26). This experiment showed that these large aggregates are indeed outside the nuclear perimeter (Figure S3A), indicating that HspB2, upon co-expression with HspB3 at a 1:1 ratio, localizes to both nuclear and cytoplasmic aggregates. Remarkably, we found that upon expression of HspB2 alone, a fraction of the Lamin B1 was sequestered inside nuclear HspB2 condensates (Figure S3A-B). In these cells, Lamin B1 still showed a continuous lining of the nuclear envelope, suggesting the integrity of the envelope was not compromised by this sequestration. In contrast with a previous study in myoblasts (18), we did not observe colocalization of HspB2 with Lamin A (Figure S3C-D), suggesting that sequestration of Lamin A in HspB2 condensates formed in myoblasts may be mediated by factors that are not expressed in HEK293 cells. In line with their liquid nature, condensates of all sizes formed upon expression of HspB2 alone appear largely circular (Figure 2C-D). In contrast, HspB2:HspB3 aggregates appear far less circular, further indicating that these structures are likely not formed through liquid-liquid phase separation.

To assess HspB2 subcellular localization, as well as the number, size, and relative intensities of HspB2 condensates and aggregates three-dimensionally, we performed high throughput confocal Z-stack imaging, followed by automated image analysis (see Figure S4 and accompanying Supplemental text for validation of the method). First, we assessed subcellular localization globally, by assessing the relative amount of HspB2-signal in the nucleus and the cytoplasm. This analysis revealed that in cells transfected with HspB2 alone, the vast majority (91%) of HspB2 resides within the nucleus (Figure 2E). Upon co-expression of a small amount of HspB3 (HspB2:HspB3 9:1), the subcellular distribution of HspB2 is unchanged (87% nuclear – Figure 2E). However, instead of being localized in nuclear condensates, HspB2 was diffusely distributed throughout the nucleus at this ratio (Figure 2A). Transfection of HspB2 and HspB3 at an equimolar ratio (HspB2:HspB3 1:1) resulted in re-distribution of HspB2 over both the nucleus (49%) and cytoplasm (51% - Figure 2E). Next, we evaluated the size of HspB2 condensates and HspB2:HspB3 aggregates, as well as the HspB2 signal intensity within both structures. Nuclear condensates that formed upon expression of HspB2 alone were on average smaller than aggregates that were observed upon transfection of both HspB2 and HspB3 (11.5 µm^3^ vs. 34.6 µm^3^, respectively - Figure 2F). Interestingly, the average pixel intensity was higher inside condensates, indicating that the HspB2 concentration in HspB2 condensates was higher compared to HspB2:HspB3 aggregates (Figure 2F).

It has been shown previously that a myopathy-associated mutation in the HspB3 gene (HspB3-R116P) prevents HspB3 from interacting with HspB2 (18). We used this mutant to assess whether a direct interaction between HspB2 and HspB3 is required for the formation of HspB2:HspB3 aggregates. As in previous experiments (Figure 2A,I), co-transfection of HspB2 with a high amount of wild-type (WT) HspB3 (9:3) caused colocalization of both proteins in amorphous aggregates (Figure S2C-D). Replacing HspB3-WT with HspB3-R116P, however, did not result in the formation of such aggregates (Figure S2C-D), but rather localization of HspB2 in nuclear condensates similar to those formed upon expression of HspB2 alone. These findings indicate that a direct interaction between HspB2 and HspB3 is required for their colocalization in aggregates. As observed previously (27), R116P-HspB3 also localized to nuclear foci, which were distinct from HspB2-WT condensates (Figure S2C-D – bottom panels), indicating that both proteins can translocate across the nuclear membrane independently, despite lacking canonical nuclear localization sequences.

Separation of the soluble and insoluble fractions of lysates prepared from cells expressing HspB2 and HspB3 at various ratios, confirms that the diffusely distributed HspB2 in the 9:1 condition forms mostly soluble complexes (Figure 2G and Supplemental Dataset 1). In contrast, in cells expressing HspB2 alone, or HspB2 and HspB3 at a 9:3 ratio, a sizeable fraction of the chaperone proteins is present in complexes that are insoluble by detergents. We quantified the data from three independent experiments and found that HspB2 is most soluble upon co-expression of an intermediate amount of HspB3 (9:1), while shifting the stoichiometry in either direction resulted in accumulation of HspB2 in the insoluble fraction (Figure 2H).

In several contexts, it has been shown that liquid droplets may transition into more solid-like aggregates (28–30). To assess whether HspB2:HspB3 aggregates form through gradual solidification of liquid droplets, we investigated the development of HspB2 condensates and aggregates over time. Briefly, we co-transfected GFP-HspB2 and untagged HspB2 (1:8) with varying amounts of HspB3, and followed the expression and localization of GFP-HspB2 by live imaging. In the absence of HspB3, HspB2 accumulated in dynamically moving, circular condensates that progressively increased in size (Figure 2I – top row and Video S3A). Upon addition of a small amount of HspB3 (HspB2:HspB3 9:1), HspB2 was uniformly distributed and its expression level gradually increased (Figure 2I – middle row and Video S3B). When further increasing the relative amount of HspB3 (HspB2:HspB3 9:3), HspB2 formed amorphous aggregates which became observable approximately 18 hours after transfection, and increased in size at later timepoints (Figure 2I – bottom row and Video S3C). Importantly, from the moment they became detectable, these structures were not circular, suggesting that they did not form through fusion and gradual solidification of liquid droplets, but rather started out as amorphous aggregates whose size steadily increased over time. Altogether, these data indicate that tight regulation of the expression of HspB2 and HspB3 is highly important for the correct subcellular distribution of these chaperone proteins.

### HspB2 condensate and HspB2:HspB3 aggregate formation is fully reversible

To probe the reversibility of HspB2 condensate and HspB2:HspB3 aggregate formation, we performed a sequential transfection experiment, in which we first expressed HspB2 and HspB3 at ratios for which condensates and aggregates are readily formed (HspB2 alone and HspB2:HspB3 1:1, respectively) and subsequently performed a second transfection, which shifted the ratio between HspB2 and HspB3 (Figure 3A). Only in the second transfection, we added GFP-tagged HspB2 at a 1:8 ratio with untagged HspB2 to distinguish between cells hit during the first and second transfection. Afterwards, we assessed the effect of shifting the HspB2:HspB3 balance on HspB2 condensates and aggregates.

**Figure 3.**
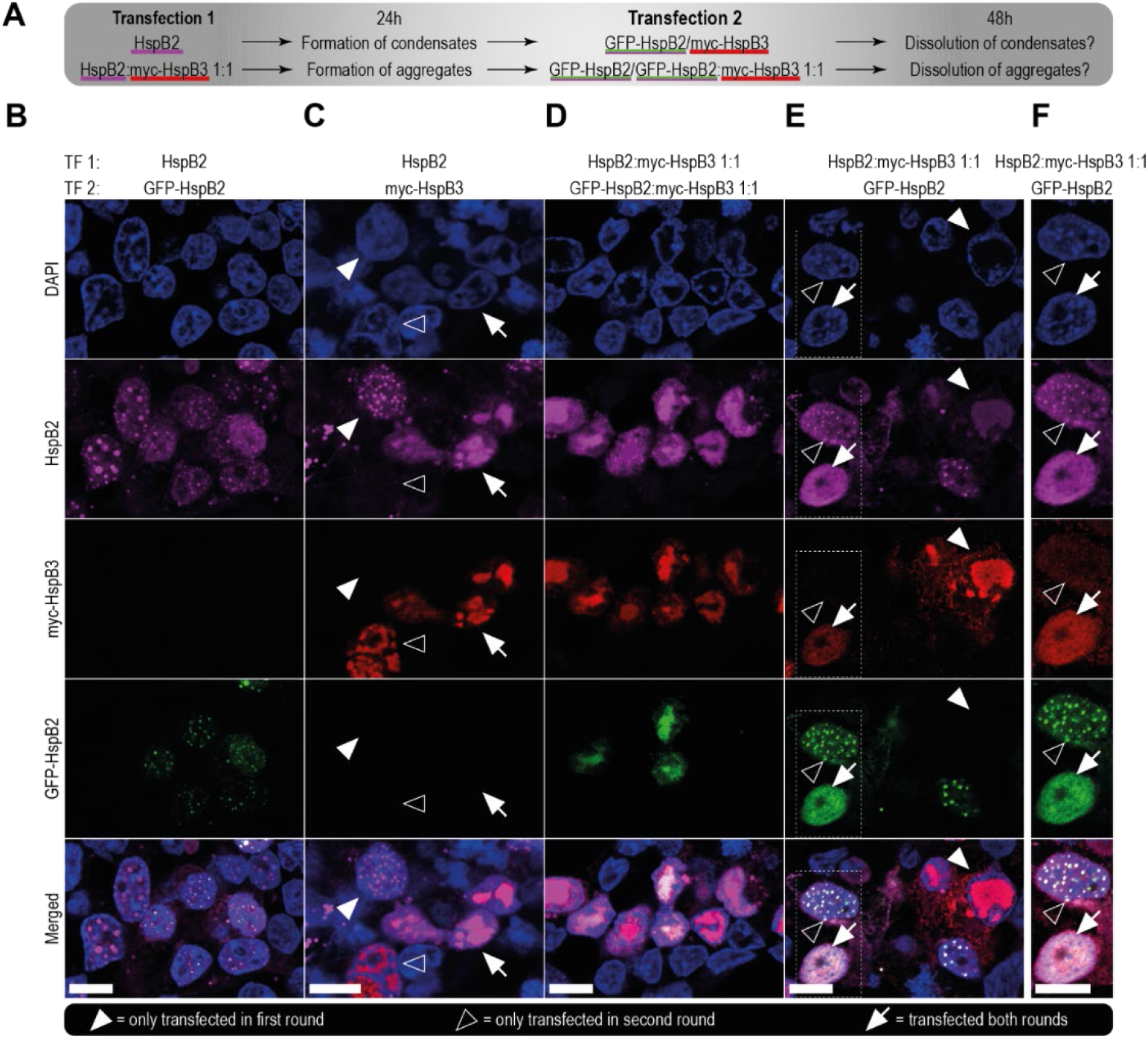
HspB2 condensates and aggregates are reversible and exchange proteins with their surroundings. **(A)** Schematic overview of the transfection schemes used in the experiment performed to generate the data presented in Figure panels (B-F). The colored underlining corresponds to the colors used for visualization of the various proteins in (B-F). **(B-E)** Confocal images depicting HEK293 cells transfected using the transfection schemes shown in (A). Closed arrowheads indicate cells hit with the first transfection (TF1), but not the second; open arrowheads indicate cells hit only in the second transfection (TF2). Arrows indicate cells that were transfected in both rounds. All scalebars are 10µm. **(F)**Magnification of the area denoted by the dashed box in (E). The myc-HspB3 signal was digitally enhanced to improve visualization of weaker signals. The scalebar represents 10µm.

In cells transfected with HspB2 alone, we readily detected nuclear HspB2 condensates (Figure 3B and closed arrowheads in Figure 3C). Subsequent transfection with myc-tagged HspB3 led to the formation of HspB3 aggregates in cells that did not express HspB2 (open arrowheads in Figure 3C), and amorphous nuclear aggregates containing both HspB2 and HspB3 in HspB2-expressing cells (arrow in Figure 3C). These findings indicate that addition of HspB3 disrupts pre-formed HspB2 condensates, resulting in the nuclear aggregation of both chaperones.

In cells transfected with HspB2- and HspB3-expressing plasmids at a 1:1 ratio, protein aggregates were readily formed (Figure 3D and closed arrowheads in Figure 3E). Surprisingly, shifting the HspB2:HspB3 balance by expressing additional HspB2 in a second transfection resulted in the dissolution of aggregates (arrow in Figure 3E, 3F). Contrastingly, cells hit only in the second transfection round (HspB2 alone) displayed the characteristic circular HspB2 condensates (open arrowheads in Figure 3E-F). These data show that HspB2 condensate and aggregate formation is reversible, suggesting that the underlying intermolecular interactions are weak and that both types of HspB2 foci are able to dynamically exchange biomolecules with the surrounding solution.

### Proximity labeling of HspB2 condensates and aggregates

Both condensates and aggregates are expected to interact with, and potentially trap specific sets of proteins, which may be linked to associated pathologies. To identify these proteins and elucidate the composition of HspB2 condensates and aggregates, we employed APEX2-mediated proximity labelling (22). In brief, plasmids encoding APEX-tagged transgenes were transfected into HEK293 cells at indicated ratios to trigger the formation of condensates and aggregates. As controls, cells transfected with plasmids encoding the APEX-tag fused to a nuclear localization signal (NLS) or nuclear export signal (NES), and cells expressing only untagged HspB2 (no APEX) were used (see Figure S5A for transfection mixes). Upon incubation with biotin-phenol and subsequent treatment with hydrogen peroxide, the APEX-tag generates short lived biotin-phenoxyl radicals which covalently bind to proximal proteins (31,32). After labelling, biotinylated proteins were enriched on streptavidin-conjugated beads and digested for analysis by mass spectrometry (Figure 4A).

**Figure 4.**
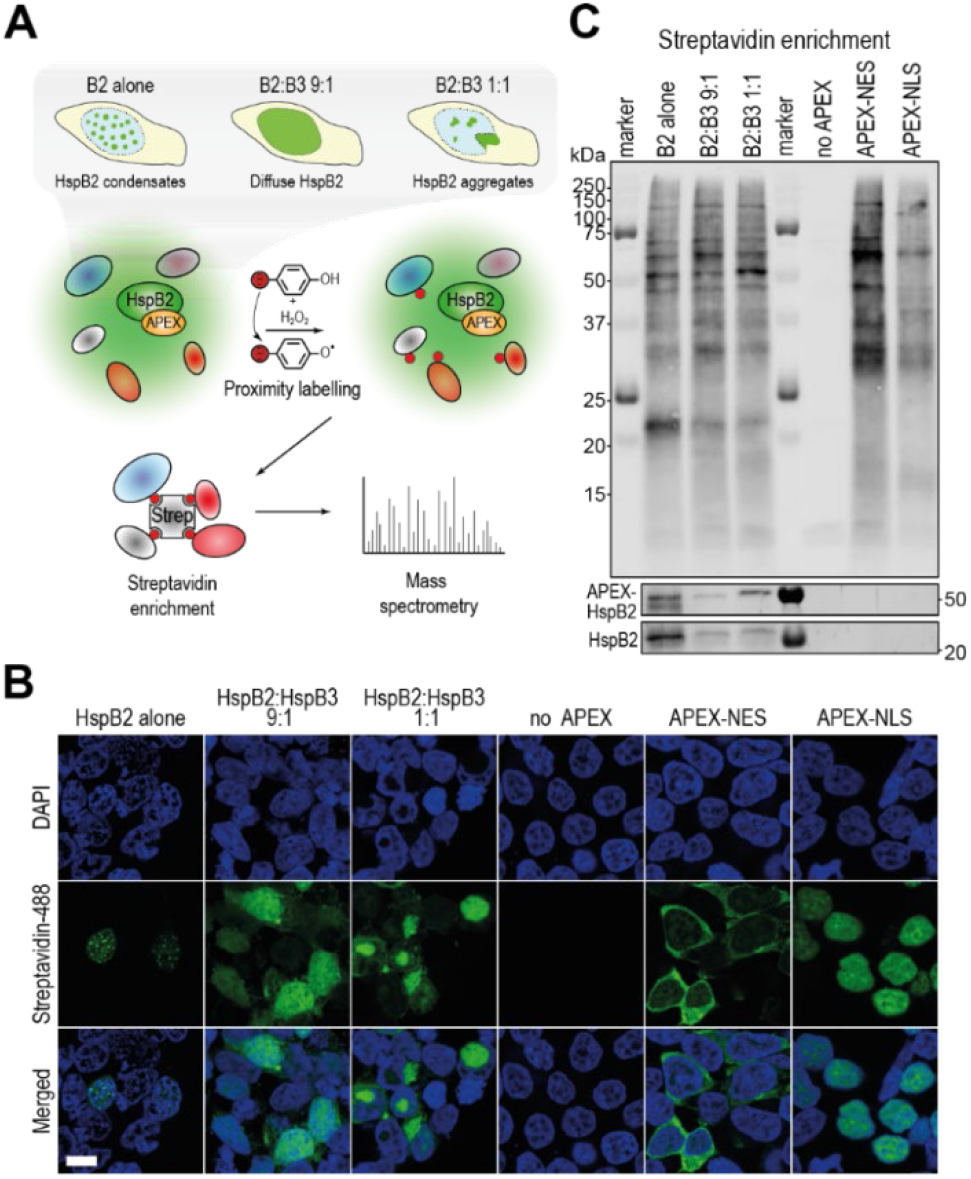
APEX-mediated proximity labelling to uncover the composition of HspB2 condensates and aggregates. **(A)** Schematic representation of the APEX-mediated proximity labelling approach used to determine the composition of HspB2 condensates and aggregates. **(B)** Confocal microscopy imaging of APEX-biotinylation activity in HEK293 cells expressing indicated transgenes. The scale bar denotes 10µm. **(C)** Streptavidin blotting of enriched proteins from lysates of cells transfected with indicated transgenes. APEX-tagged and untagged HspB2 was detected using an antibody targeting HspB2.

We first evaluated the specificity of APEX-proximity labelling by visualizing biotinylation with an Alexa488-fluorophore coupled streptavidin. For all conditions, the biotinylation patterns mirrored those observed previously when using untagged or GFP-tagged constructs: cells expressing HspB2 alone displayed biotinylation of small circular nuclear condensates, expression of HspB2 and HspB3 at a 9:1 ratio resulted in diffuse labelling throughout the nucleus, and cells transfected with HspB2 and HspB3 in a 1:1 ratio display labelling of large cytoplasmic and nuclear aggregates (Figure 4B). As expected, the APEX-NES and APEX-NLS controls show cytoplasmic and nuclear biotinylation, respectively, while no labelling is observed in cells lacking an APEX-construct (no APEX). We next performed streptavidin enrichment on lysates from cells expressing the various constructs mentioned above, and assessed biotinylation patterns and enrichment efficiency through western blotting (Figure 4C and Figure S5B).

### Unbound nuclear proteins can freely shuttle in and out of HspB2 condensates

Proteins biotinylated in HspB2 condensates and aggregates were purified on streptavidin beads, and digested to peptides which were measured by mass spectrometry. We first assessed proteins enriched in, and depleted from nuclear HspB2 condensates which formed after the expression of HspB2 alone. As these condensates are virtually exclusively nuclear (Figure 2E), we calculated enrichments relative to the APEX-NLS control. As expected, HspB2 is dramatically enriched (>26,000-fold; Figure 5A), indicating efficient auto-biotinylation. In line with previous studies (33,34), we found enrichments of HspB1 (22-fold) and BAG3 (6-fold) in HspB2 condensates, indicating that our method is able to identify known interactions. In addition, we uncover previously unknown interactions with various proteins including LGALS7 (~78-fold), and MEA1 (~26-fold). For gene ontology (GO)-term enrichment analysis, we selected the top 5% enriched factors with a *P*-value < 0.01 (blue in Figure 5A, listed in Table S1A). This analysis uncovered an enrichment of components of the HAUS complex (Figure 5B); a complex that localizes to the centrosome during in mitotic spindle assembly.

**Figure 5.**
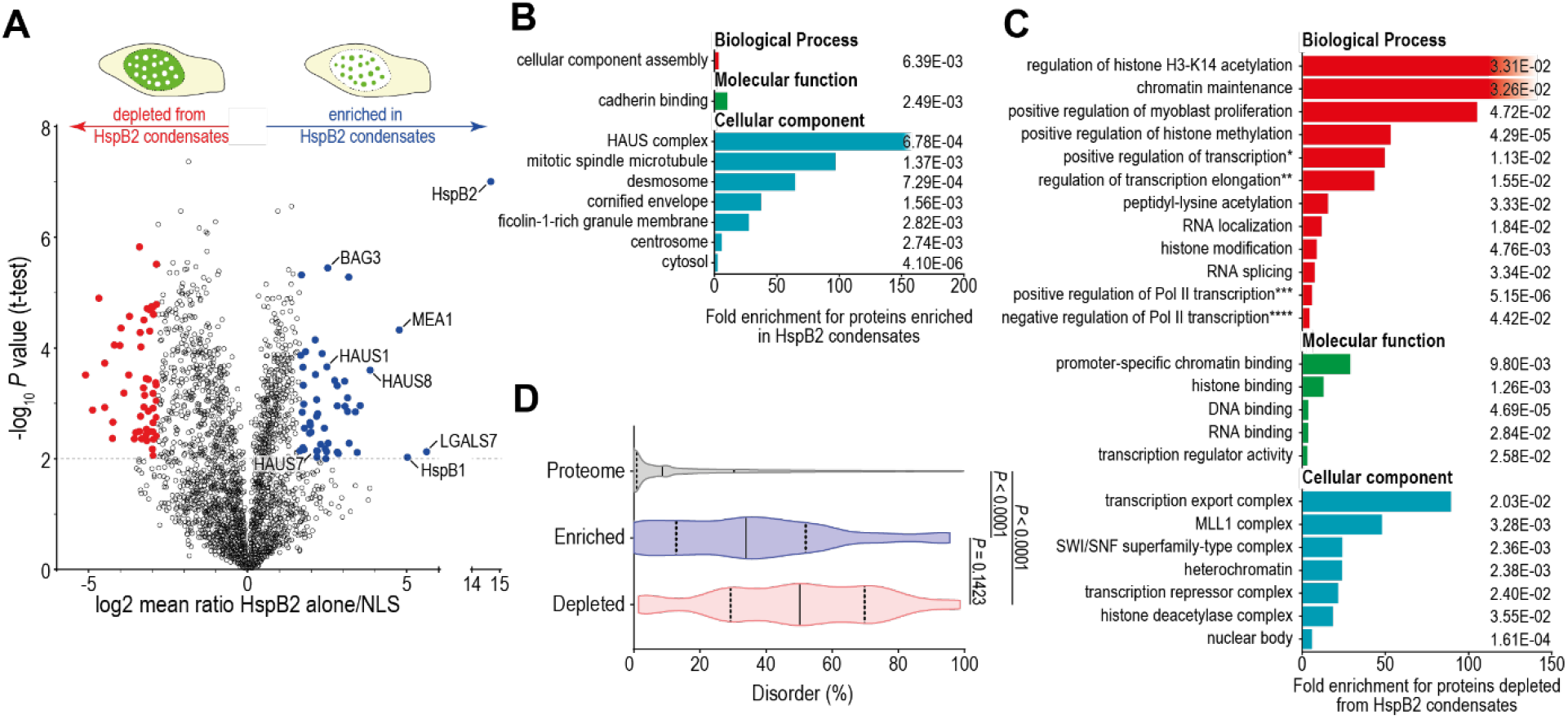
DNA-associated proteins are depleted from nuclear HspB2 condensates. **(A)** Volcano plot depicting the mean ratio of proximal proteins detected upon APEX-labelling in cells expressing APEX-tagged HspB2 and APEX-NLS. The top 5% proteins enriched in HspB2 condensates are depicted in blue, red dots indicate the 5% proteins most depleted from HspB2 condensates. **(B)** GO-term enrichment analysis for factors enriched in HspB2 condensates (blue in (A)), as determined using gene ontology analysis. Only significantly enriched terms are shown, bars depict the fold enrichment over background (human proteome), and the numbers indicate the FDR as determined on http://geneontology.org/. **(C)** Same as (B) but for factors depleted from HspB2 condensates (red in (A)). Four terms are abbreviated due to space constraints (marked with asterisks). Full terms are: * positive regulation of DNA-templated transcription, elongation; ** regulation of transcription elongation from RNA-Pol II promoter, *** positive regulation of transcription by RNA-Pol II; **** negative regulation of transcription by RNA-Pol II. **(D)** Violin plots depicting the percentage distribution of disordered residues per protein in the human proteome (grey; n=61403), HspB2 condensate components (blue; n=49), and proteins depleted from HspB2 condensates (red; n=49) as predicted by the IUPred algorithm. Solid and dashed lines indicate the median and quartiles, respectively. A Kruskal-Wallis test followed by Dunn’s multiple comparisons test was used to determine statistical significance.

To analyze proteins that were depleted from HspB2 condensates, we next assessed GO-term enrichment for the top 5% depleted factors with a *P*-value < 0.01 (red in Figure 5A, listed in Table S1B). This analysis uncovered that proteins bound to chromatin, histones and DNA are significantly depleted from HspB2 condensates (Figure 5C), suggesting that these proteins are unavailable for shuttling into the condensates because of their association with the genome. In line with this, proteins enriched in HspB2 condensates are depleted for the GO-terms chromatin-binding, histone binding and DNA binding (n=0, n=0, and n=2 out of 48 proteins, respectively).

As intrinsic protein disorder is considered to be a major contributing factor in condensate formation (35), we subsequently compared the amount of disorder in proteins enriched in, and depleted from HspB2 condensates. As expected, proteins enriched in condensates are significantly more disordered compared to the amount of disorder found across the human proteome (median disorder 33.9% vs. 8.6%, respectively; Figure 5D). Intriguingly however, proteins depleted from condensates displayed an even larger degree of disorder (50.2%). This can be explained by the fact that, in general, nuclear proteins display a larger degree of intrinsic disorder (25.2% - Figure S6A) compared to cytoplasmic proteins (11.0%) or proteins that shuttle between the two compartments (17.0%). Furthermore, the high degree of disorder in proteins depleted from HspB2 condensates is in line with findings that disorder is particularly enriched in transcription factors and chromatin remodelers (36–38).

While it is interesting to identify specific proteins that were enriched or depleted from condensates, perhaps the most striking result from our proximity labeling analysis is the fact that the vast majority of proteins are neither enriched nor depleted. We found that for HspB2 alone/NLS, 97.4% of all identified proteins had log2 enrichment scores between −2.89 (*e*^−2^) and 2.89 (*e*^*2*^), corresponding to a partitioning free energy difference of less than 2 k_B_T (Figure S6B). This suggests that most of the detected nuclear proteins are able to shuttle freely in and out of HspB2 condensates, and have a negligible preference for being inside the condensate over the nucleoplasm, or vice versa. If they would have had a strongly unfavorable interaction with condensate components, they would hardly enter the condensates and we would expect these proteins to be strongly depleted in Figure 5A. On the other hand, if they would have a strongly favorable interaction with condensate components, they would be unlikely to leave the condensates and we would expect these proteins to be enriched. Our findings suggest that the vast majority of unbound nuclear proteins may shuttle freely in and out of condensates, without getting trapped.

### Autophagy factors are recruited to HspB2:HspB3 aggregates

We next set out to elucidate the composition of the amorphous aggregates formed in cells transfected with HspB2 and HspB3 at an equimolar ratio. As these aggregates are equally present in the nucleus and cytoplasm (Figure 2E), we calculated enrichments relative to both APEX-NES and APEX-NLS controls. HspB2 and HspB3 were strongly enriched (13,000-fold (NES)/22,000-fold (NLS) and 12,000-fold (NES)/15,000-fold (NLS), respectively), indicating a very strong interaction between the two chaperone proteins (Figure 6A-B). As seen for the HspB2 condensates, known interactors BAG3 (11-fold (NES)/23-fold (NLS)) and HspB1 (8-fold (NES)/43-fold (NLS)) were also enriched in HspB2:HspB3 aggregates. Similarly, HspB1 and BAG3 were enriched upon HspB2 proximity labelling in cells where HspB2 was diffusely distributed throughout the nucleus (HspB2:HspB3 9:1), indicating these interactions are independent of the subcellular localization or phase behavior of HspB2.

**Figure 6.**
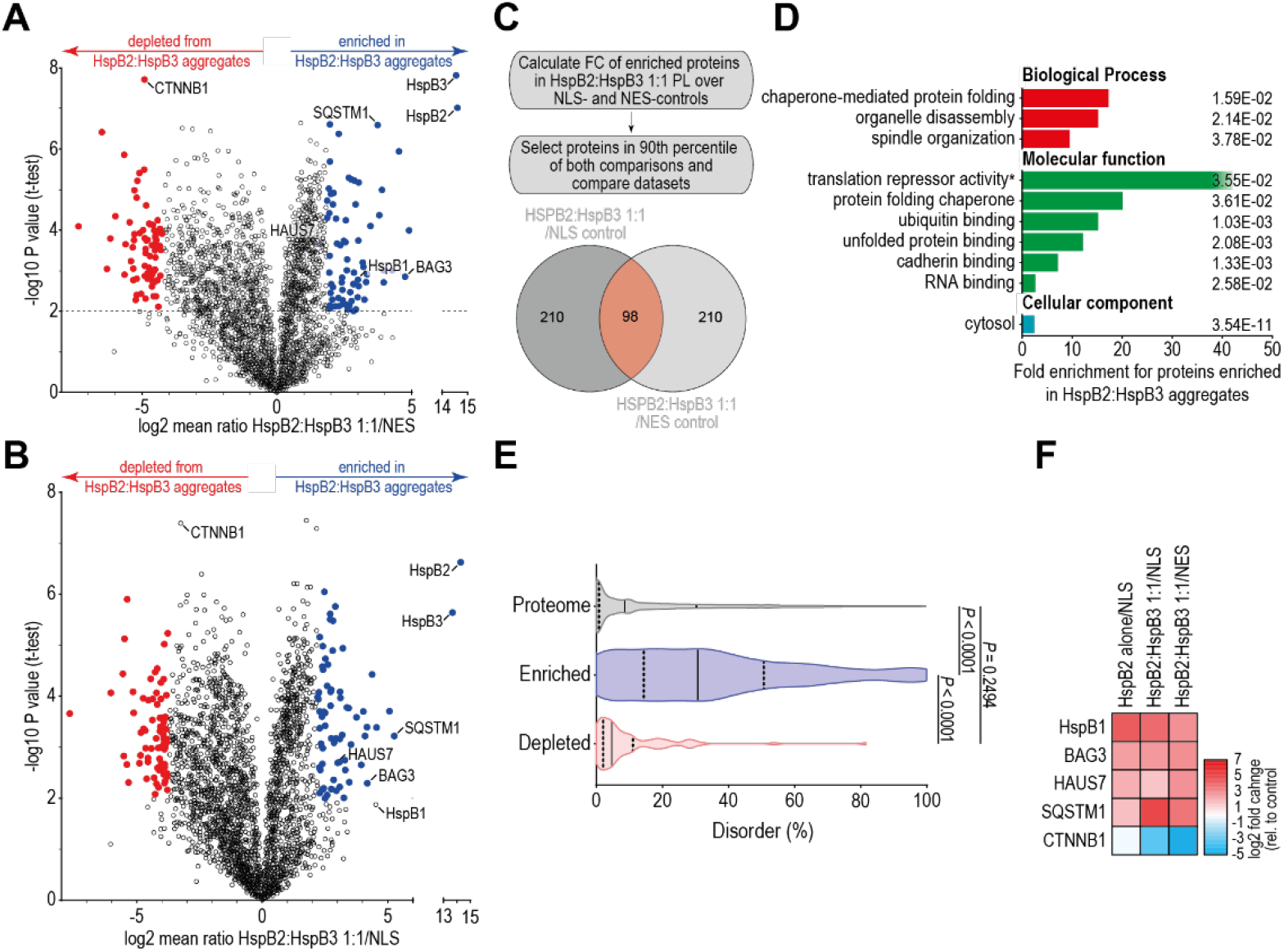
Autophagy factors are recruited to HspB2:HspB3 aggregates. **(A-B)** Enrichment of biotinylated proteins in HspB2:HspB3 proximity labelling relative to APEX-NES (A) and APEX-NLS (B) controls. The 5% most enriched and depleted proteins with a *P*-value <0.01 are shown in blue and red, respectively. **(C)** Overview of the method used to select a high-confidence set of proteins associated with HspB2:HspB3 aggregates by comparing against two different control datasets. **(D)** Gene onthology (GO)-term enrichment analysis of the proteins identified in (C) as determined using the http://geneontology.org/ webtool. Only significant term are shown, numbers indicate the FDR. * GO-term abbreviated, full term: ‘translation repressor activity, mRNA regulatory element binding’. **(E)** Violin plots displaying the distribution of protein disorder in the human proteome (grey; n=61403), proteins enriched (blue; n=98) and depleted (red; n=87) in HspB2:HspB3 aggregates as predicted by IUPred. The median and quartiles are indicated by solid and dashed lines, respectively. To test for significance, a Kruskal-Wallis test followed by Dunn’s test for multiple comparisons was used. **(F)** Summary of the log2 fold changes of the five proteins used for validation in our HspB2 proximity labelling datasets, relative to indicated relevant controls.

We identified the top 10% enriched proteins relative to both APEX-NES and APEX-NLS controls, and selected those proteins (n=98) present in both comparisons for GO-term enrichment analysis (Figure 6C, listed in Table S1C). This analysis revealed an enrichment of factors associated with chaperone-assisted protein folding (Figure 6D). In line with this, the amount of disorder in proteins enriched in HspB2:HspB3 aggregates is dramatically higher compared to the proteome (median disorder 30.7% vs. 8.6%, respectively; Figure 6E), suggesting that many disordered and/or misfolded proteins are associated with these aggregates. As seen for HspB2 condensates, proteins involved in the organization of the mitotic spindle are enriched in HspB2:HspB3 aggregates. Moreover, enrichment of both RNA binding proteins and proteins involved in translation repression suggests that HspB2:HspB3 aggregates may influence RNA processes, analogous to stress granules. Lastly, factors involved in organelle disassembly (autophagy) such as SQSTM1 (also known as p62) and ATG3 are strongly enriched, suggesting that the cell is attempting to clear these aggregates.

Interestingly, significantly more proteins were enriched in HspB2:HspB3 aggregates, as compared to HspB2 condensates (1.2% vs. 0.6%, respectively - Figure S6B), indicating that a sizeable group of proteins is effectively trapped within HspB2:HspB3 aggregates. Similarly, an even larger subset of proteins is depleted from aggregates as compared to condensates (8.1% vs. 2.0%), indicating HspB2:HspB3 aggregates may constitute a less favorable environment for a larger group of proteins.

### Proximity labelling of another type of sHSP aggregate reveals enrichment of common factors

To enable comparison with another aggregation prone sHSP, we concurrently performed proximity labelling of HspB5-STD/RG, a myopathy-related HspB5 mutant which contains an arginine to glycine substitution at position 120 (R120G), as well as three serine to aspartic acid phosphomimicking mutations at positions 19, 45 and 59 (S19D, S45D, S59D). Prior work has shown that this mutant accumulates in cytoplasmic aggregates (42,43), similar to those seen for HspB2:HspB3. Mapping the composition of two types of sHSP-containing aggregates allows the identification of potentially common factors recruited to such structures.

Proximity labelling in cells transfected with APEX-tagged HspB5-WT and -STD/RG constructs resulted in distinct biotinylation patterns (Figure S7A). Biotinylation in HspB5-WT cells localized diffusely in the cytoplasm, while in HspB5-STD/RG cells, amorphous cytoplasmic aggregates were observed (Figure S7B). As HspB5-STD/RG is exclusively cytoplasmic, we determined enrichment of biotinylated proteins relative to the APEX-NES control (Figure S7C, enriched and depleted proteins are listed in Table S1D-E). HspB5 was enriched very strongly (1200-fold), indicating that proximity labelling was successful. Strikingly, numerous proteins that were found in HspB2:HspB3 aggregates were also enriched in HspB5-STD/RG proximity labelling, including SQSTM1, BAG3, and HspB1 (Figure S7C). This prompted us to overlap the sets of enriched proteins in HspB2:HspB3- and HspB5-STD/RG aggregates to identify common factors. To this end, we selected the top 5% enriched proteins with a *P*-value <0.01 from both datasets relative to APEX-NES control and selected common factors (n=13; Figure S7D). Among these common factors were several proteins known to be involved in protein folding, ubiquitination and autophagy (Figure S7E). These findings suggest that a common machinery is recruited to clear aggregates centered around different small heat shock proteins.

### Validation of protein components in HspB2 condensates and HspB2:HspB3 aggregates

To validate our proximity labelling proteomics datasets, we selected proteins with distinct enrichment patterns across the various conditions. HspB1, BAG3, and HAUS7 were equally enriched in HspB2 condensates and HspB2:HspB3 aggregates, while SQSTM1 was predominantly enriched in HspB2 aggregates, and CTNNB1 was depleted from both structures (Figure 6F). HspB1, BAG3, and SQSTM1 were also enriched in HspB5-STD/RG aggregates, while HAUS7 and CTNNB1 were depleted (Figure S7F). A schematic representation of domain composition, charge distribution and predicted disorder of these proteins is shown in Figure S6C.

We first assessed colocalization of candidate proteins with HspB2 condensates. Both HspB1 and BAG3 have been shown to reside in the cytoplasm under physiological conditions, and translocate to the nucleus in response to heat shock (44,45). Yet, in our experiments, the proteins are present in both compartments under physiological conditions (Supplemental Dataset 2). Upon HspB2 expression, HspB1, BAG3, and HAUS7 all colocalize with nuclear HspB2 condensates (Figure 7A-C), confirming our proximity labelling dataset. In line with earlier work (46), SQSTM1 localized to cytoplasmic and nuclear foci, yet, these do not colocalize with HspB2 condensates (Figure 7A-C). Previous work showed the *O*-GlcNAcylation-dependent translocation of CTNNB1 to the plasma membrane (47). In line with this, we observe CTNBB1-staining at the cell membrane, which therefore does not overlap with the nuclear HspB2-signal (Figure 7A-C). HspB1, BAG3, and SQSTM1 all colocalized with HspB2:HspB3 and HspB5-STD/RG aggregates, while HAUS7 colocalized only with HspB2:HspB3 aggregates (Figure 7D, Figure S7G). Lastly, CTNNB1 is depleted from both aggregates (Figure 7D, Figure S7G). Altogether, the colocalization imaging experiments presented here mirror the enrichment patterns observed in our proximity labelling experiments, thereby validating our proteomics datasets.

**Figure 7.**
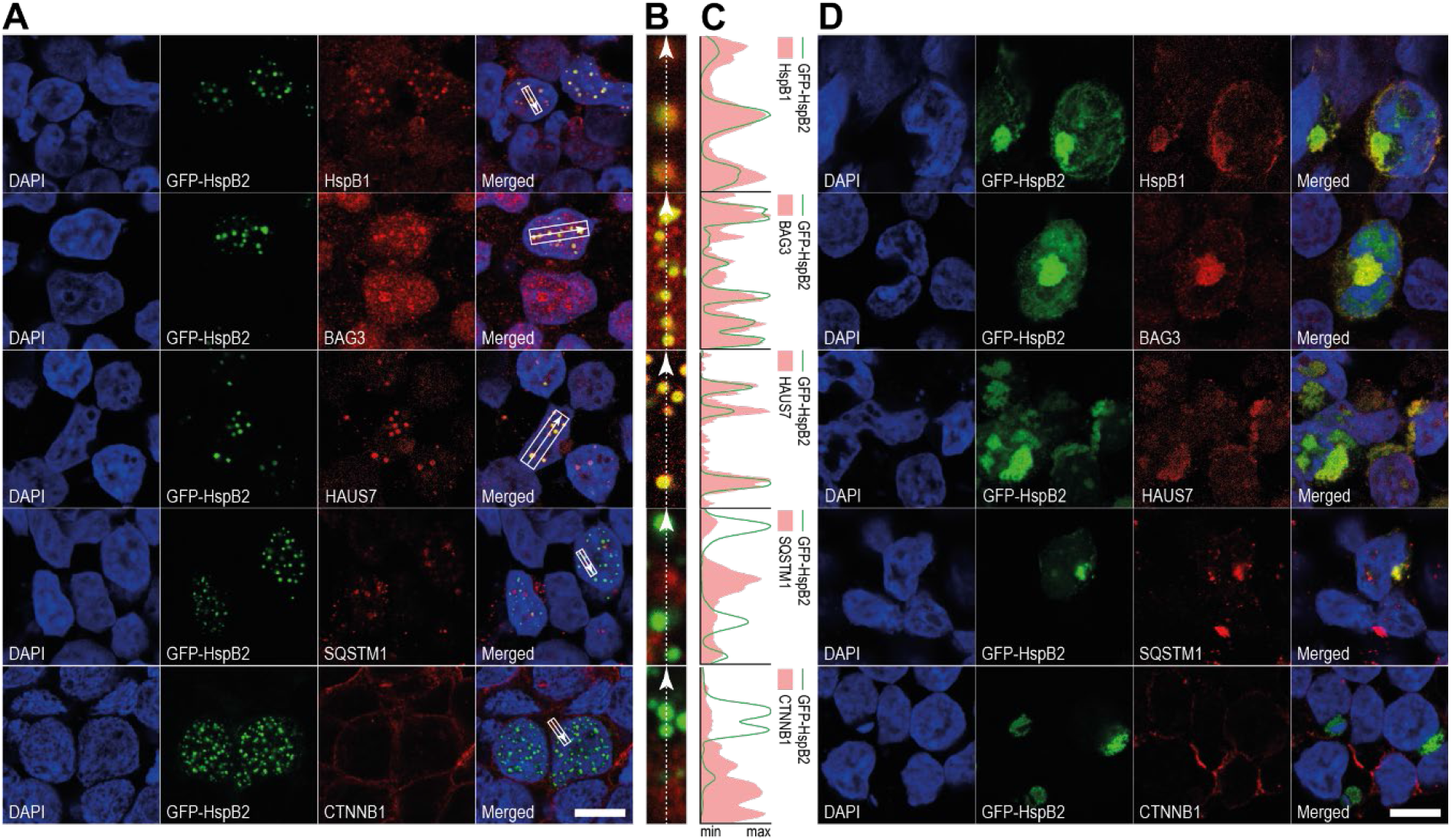
Validation of proximity labelling datasets through colocalization studies. **(A)** Confocal microscopy images of nuclear condensates in HEK293 cells expressing HspB2 alone, co-stained for indicated interactors. The rectangular inset denotes the area of the image shown enlarged in (B). The scalebar denotes 10µm (same for (D)). **(B)** Magnification of the inset shown in (A). The arrow indicates the line and direction along which signal intensity was determined in (C). **(C)** Histogram depicting signal intensities of GFP-HspB2 (green lines) and indicated candidates (red shading) along the arrows shown in (B). **(D)** Confocal images of protein aggregates in HEK293 cells transfected with HspB2 and HspB3 at a 1:1 ratio, co-stained with antibodies targeting indicated interactors.

## DISCUSSION

Small heat shock proteins (sHSPs) are the most ubiquitous class of molecular chaperones, and are conserved across all branches of life (48). A key characteristic of sHSPs is that they assemble into polydisperse, homo- and hetero-oligomeric assemblies of varying conformations and sizes (14,15). It is widely believed that smaller oligomers are more active as chaperones, while larger oligomers likely function as storage complexes (49). Interestingly, the chaperone activity of sHSPs can be dynamically regulated by shifting the equilibrium between different sHSPs (21). To investigate the effects of shifting the balance between components of sHSP hetero-oligomers inside living cells, we focused on HspB2 and HspB3, two chaperones with overlapping tissue expression patterns that have been shown to directly interact (18–20) and co-assemble at a well-defined 3:1 ratio in vitro (20).

In this study, we explored the effect of changing HspB2:HspB3 expression ratios on the subcellular localization and phase behavior of these proteins inside living cells. To ensure full control over relative HspB2 and HspB3 expression levels without the need for suboptimal knockdown or laborious knockout approaches, we used HEK293 cells, as they do not express HspB2 and HspB3 endogenously. This experimental set-up therefore provided a clean slate for the study of the effects of stoichiometric shifts in the HspB2:HspB3 balance. We assessed the effects of altering intracellular HspB2:HspB3 ratios by transfecting plasmid DNA encoding these chaperones at various ratios. Guided by the findings in (50), we ensured that the amount of DNA transfected did not saturate the cellular transcription machinery. Moreover, expression of (GFP-)HspB2 and HspB3 was driven by the same cytomegalovirus (CMV) immediate-early enhancer and promoter to minimize potential variations in transcription efficiency, thus ensuring that HspB2:HspB3 DNA ratios correlated linearly with the relative proteins levels.

Numerous studies have shown that homo- and hetero-oligomeric sHSP assemblies dramatically differ in their chaperone activity (21,51,52). For instance, HspB2:HspB3 hetero-oligomers have poor chaperone activity (20), while HspB2 and HspB3 homo-oligomers display moderate chaperone activities, as compared to HspB5 (53–55). In contrast, hetero-oligomers of HspB4 and HspB5 outperform homo-oligomers consisting of either of the two proteins in chaperone activity assays (56), and hetero-oligomerization with HspB6 increases chaperone activity of HspB5 (21). While such insights are invaluable, these studies rely on in vitro incubation of purified chaperones with client proteins, thus ignoring the complexity of the cellular environment in which these proteins naturally operate. Our experimental approach of controllable sHSP expression in HEK293 cells can be used to investigate sHSP oligomer assembly inside living cells.

HspB2 readily phase separates into nuclear droplets in a concentration-dependent manner in HEK293 cells (this study), as well as in differentiating myoblasts and HeLa cells (18), suggesting that condensate formation is an intrinsic feature of HspB2. Yet, the latter study also reported the accumulation of overexpressed HspB2 in large amorphous intranuclear compartments, while in our study, overexpression of HspB2 by itself resulted in either diffuse distribution of the protein (at low concentrations) or condensation into liquid droplets (at higher concentrations). Moreover, Lamin A is sequestered inside HspB2 compartments in differentiating myoblasts (18), but not in HspB2 condensates in HEK293 cells. These findings suggest that HspB2 phase behavior is, in part, cell type specific, which may result from variations in the expression patterns of interacting proteins (such as Lamin A) and other biomolecules, as has been shown previously for other interaction networks (57–59). Other sHSPs have been found in condensates or condensate-like structures. For instance, HspB1 and HspB8 locate to stress granules upon proteotoxic stress (60), and both HspB7 and pseudophosphorylated HspB5 are found in nuclear granules (44,61). However, condensate formation induced solely by overexpression is a unique feature of HspB2. The reason why HspB2 is able to form condensates on its own remains unclear, but may be related to another unique feature of HspB2, namely its ability to form HspB2/HspB3 tetramers with an anomalous 3:1 ratio. The formation of this unusual tetrameric conformation may require attractive homotypic HspB2-HspB2 interactions that could also underlie condensate formation in the absence of HspB3.

Phase separated condensates often serve to concentrate biochemical reactions within cells (62,63). It is therefore tempting to speculate that HspB2 condensates may promote the concentration of chaperones and clients to promote chaperone-guided protein refolding. Arguing against this hypothesis however, is the fact that HspB2 condensates are not particularly enriched for disordered proteins and that traditional chaperone clients such as FUS and α-synuclein are not enriched in HspB2 condensates. It is more likely that HspB2 condensates serve as a way to prevent potential toxic effects of excessive HspB2 levels in absence of its partner HspB3. In addition, HspB2 condensates may serve as nuclear chaperone storage pools, from which HspB2 can rapidly be released in times of stress.

Co-expression of an intermediate amount of HspB3 (HspB2:HspB3 9:1) prevented HspB2 phase separation, while addition of a larger amount of HspB3 (9:3 or 9:9) led to the formation of large cytoplasmic and nuclear aggregates. These findings illustrate the dramatic effects of stoichiometric shifts in hetero-oligomer composition, and underscore the importance of maintaining the correct expression balance for sHSP physiological functioning. Remarkably, we observed no obvious detrimental effects on cell survival and proliferation, suggesting that both structures are tolerated by the cells. HspB2 condensates can be dissolved by expression of HspB3 at a later stage, showing that HspB2 condensates are reversible and exchange biomolecules with their surroundings. To our surprise, we also found that HspB2:HspB3 aggregates can be dissolved by shifting the ratio towards HspB2, indicating that formation of these solid-like structures is also reversible, at least in the timeframe we assessed. These results show that both condensates and aggregates are likely formed by transient, weak interactions with other cellular proteins. Whether the observed structures may have longer-term toxic effects remains to be seen.

APEX-mediated proximity labelling of HspB2 condensates revealed that only few of the detected proteins are strongly enriched in, or depleted from HspB2 condensates, suggesting that most proteins are able to freely diffuse in and out of these droplets, without getting trapped. In comparison, substantially more proteins are enriched in, and depleted from solid-like HspB2:HspB3 and HspB5-STD/RG aggregates, indicating subsets of proteins are specifically trapped inside these structures. Interestingly, a specific group of proteins is enriched in both HspB2:HspB3 and HspB5-STD/RG aggregates. This group includes proteins involved in ubiquitination and autophagy including SQSTM1/p62, and may therefore represent a common molecular machinery aimed at clearing sHSP aggregates. SQSTM1/p62 accumulation has previously been observed in HspB5-STD/RG aggregates (64), and is often interpreted as a marker for defects in autophagic degradation (65,66). Whether autophagy of the sHSP aggregates observed here is effective, or inhibited because of entrapment of these factors within aggregates remains to be explored.

In differentiating myoblasts, HspB2 subcellular localization is highly heterogeneous, ranging from nuclear condensates and diffuse distribution, to amorphous compartments (18). This dramatic phenotypical variation likely stems from cell-to-cell variation in the differentiation stage and related HspB2:HspB3 expression levels. The HEK293 cell platform we developed here allows precise control over relative sHSP expression levels, enabling the study of sHSP structures with uniform biophysical properties and subcellular localization phenotypes. As such, this approach allowed us to map the composition of HspB2 condensates and HspB2:HspB3 aggregates, as well as HspB5-STD/RG aggregates. We anticipate that our approach can be used for the study of biophysical properties and composition of other sHSP structures, as well as the investigation of the effects of disease-related sHSP mutants (67). Our findings may help shed light on the origins of phenotypes associated with mutations in sHSPs that cause inclusion bodies, and improve our understanding of the functions and mechanisms of sHSPs.

## ACKNOWLEDGEMENTS

We thank colleagues from the departments of Biomolecular Chemistry and Physical Organic Chemistry for fruitful discussions. We thank dr. Wolf of the department of Cell Biology, RadboudUMC Nijmegen for kindly providing the Lamin A and Lamin B1 antibodies. We acknowledge dr. Jelle Postma, for his assistance in microscopic imaging. This work was financially supported by the Institute for Molecules and Materials (Radboud University). E.S. acknowledges the Netherlands Organization for Scientific Research (NWO – grant number: 193.089) for funding.

## AUTHOR CONTIBUTIONS

Conceptualization, J.J., E.S., W.B., K.M.B.; Methodology, J.J.; Software, B.v.S; Validation, J.J., W.V.E.; Formal Analysis, J.J., B.v.S., P.W.T.C.J.; Investigation, J.J., W.V.E., M.E., Resources, M.V., W.B., K.M.B., E.S.; Writing – Original Draft, J.J., E.S.; Writing – Review & Editing, J.J., E.S., W.B.; Visualization, J.J.; Supervision, M.V., W.B., K.M.B., E.S., J.J.; Funding Acquisition, W.B., K.M.B., E.S.

## CONFLICT OF INTERESTS

The authors declare that they have no conflict of interest.

## MATERIALS AND METHODS

### Analysis of sHSP expression

For the generation of the sHSP expression heatmaps depicted in Figure S1, RNA and protein expression data was extracted from the Human Protein Atlas (www.proteinatlas.org (68)). The extracted RNA-expression data is a consensus dataset derived from the RNA-seq dataset generated internally by the Human Protein Atlas (HPA), RNA-seq data from the Genotype-Tissue Expression (GTEx) project, and CAGE data from the FANTOM5 project. Expression values from the three datasets were normalized to generate the Normalized eXpression (NX) values depicted in Figure S1A-B. For the tissue enrichment analysis, a pseudocount of 0.1 was added to all NX-values. This tissue-specific expression values were subsequently divided by the average sHSP expression value across all tissues as a measure of tissue-specific enrichment. The data was then log2 transformed to produce the heatmap depicted in Figure S1C. Protein expression data is derived from immunohistochemically stained tissues, using four levels of expression (high, medium, low, and not detected).

### Protein disorder prediction

To predict intrinsically disordered regions within HspB2 and candidate proteins, we used IUPred 3 (https://iupred3.elte.hu/ (41)), using default settings. The percentage of disorder for groups of proteins identified by mass spectrometry, as well as for the entire human proteome was extracted from the Disorder Atlas (https://disorderatlas.med.umich.edu/index) (69,70), which makes use of the IUPred prediction algorithm (71,72).

### Culturing and transfecting HEK293 cells

HEK293 cells were maintained in DMEM supplemented with 10% fetal bovine serum (Sigma, #F0804) and 1% Penicillin/Streptoymycin (HyClone, SV30010) at 37°C and 5% CO_2_. Cells were detached using Trypsin/EDTA (0.25%/0.02%) and passaged two times per week at a 1:10 to 1:20 ratio, depending on confluency.

For transfection of HEK293 cells seeded into a single well of a 24-wells plate (culture volume 500 µL), 500 ng plasmid DNA was combined with 1.5 µL polyethyleneimine (PEI, 1 mg/mL, Sigma, #p3143) in 50 µL Optimem (Gibco, #31985070). The transfection mixture was incubated at room temperature for 30 minutes before being added dropwise to the cells. For larger culture volumes, the amount of plasmid DNA, PEI, and Optimem was increased proportionally.

### Live cell imaging

All live cell imaging was performed at 37 °C on the Leica DMi8 widefield microscope in Cellvis glass bottom dishes (cat# D60-30-1.5-N). During live imaging experiments, HEPES was added to the culture medium to an end concentration of 25 mM to control fluctuations in pH in the absence of CO_2_. Thunder computational clearing and widefield deconvolution was applied to live imaging data for background subtraction. For Videos S1-2, bleach correction was performed in Fiji (Image>Adjust>Bleach Correction>Histogram Matching) to correct for signal decay due to high scanning frequency.

### Immunofluorescence analysis

Coverslips were sterilized with 100% ethanol prior to coating with poly-L-lysine (0.1mg/mL) for 5 minutes at room temperature. After thorough rinsing with milli-Q, coverslips were allowed to dry for at least two hours before seeding HEK293 cells as described above. Approximately 16 hours after seeding, cells were transfected as described above. 24 hours after transfection, cells were fixed in 4% paraformaldehyde (PFA; Fluka-#47629), supplemented with 0.1% SDS, 0.5% sodium deoxycholate and 1% Triton. After fixation, cells were washed three times with PBS before incubating with 200 µL ice-cold acetone for 5 minutes. After two washes with PBS, cells were either stored under PBS-Glycine (10mM) at 4°C or directly stained as below.

Subsequently, cells were treated with 0.25% Triton in PBS for 20 minutes. Cells were subsequently washed with PBS and incubated in blocking buffer (3% BSA, 0.1% Triton, 10% normal goat serum in PBS) supplemented with 100 mM glycine for 30 minutes. After blocking, cells were incubated with primary antibodies (all at 1:100 dilution) in blocking buffer for 1 hour at room temperature. Primary antibodies used in immunofluorescence experiments were rabbit-anti-HspB2 (generated in our laboratory (73,74)), mouse-anti-myc (Thermo Fisher Scientific Cat# MA1-980, RRID:AB_558470), mouse-anti-Lamin A (Nordic-MUbio, cat# 1101P), rabbit-anti-Lamin B1 (Proteintech, cat# 12987-1-AP, RRID:AB_2136290), mouse-anti-SQSTM1 (Santa Cruz Biotechnology, cat# sc-28359, RRID:AB_628279), mouse-anti-BAG3 (Santa Cruz Biotechnology, cat# sc-136467, RRID:AB_10647772), rabbit-anti-HspB1 (Enzo Life Sciences, cat# ADI-SPA-803, RRID:AB_10615084), mouse-anti-HAUS7 (Santa Cruz Biotechnology, cat# sc-393259), mouse-anti-CTNNB1 (Santa Cruz Biotechnology, cat# sc-7963, RRID:AB_626807). After incubation with primary antibodies, coverslips were washed three times using 0.05% Tween in PBS for 5 minutes before incubation with secondary antibodies (all at 1:100 dilution) in blocking buffer for 1 hour at room temperature. Secondary antibodies used in immunofluorescence experiments were goat-anti-mouse Alexa 568 (Thermo Fisher Scientific Cat# A-11004, RRID:AB_2534072) and goat-anti-rabbit Alexa647 (Thermo Fisher Scientific Cat# A27040, RRID:AB_2536101). Where indicated, biotinylated proteins were visualized using a streptavidin-alexa488 conjugate (Invitrogen, #S32354). Subsequently, cells were washed twice with 0.05% Tween in PBS and twice with PBS before nuclei were counterstained with 1 µg/mL DAPI in PBS for 5 minutes at room temperature. Afterwards, cells were rinsed twice with PBS, once with PBS-milliQ 1:1, and three times with milliQ before mounting in Mowiol (Sigma, #81381). After overnight hardening of the mounting medium at room temperature, microscopy samples were stored at 4°C.

### Imaging of fixed cells

All images of fixed material were generated using the Leica SP8 confocal microscope, except for those shown in Figure 2G, which were made using the Zeiss Axio Imager fluorescence microscope. The histograms depicting signal intensity in Figures 2B, 7C, S2B, S3B, and S3D were generated using Fiji (75). Intensity values were optimized using the minimal and maximal intensity values as borders for each channel separately, after which smoothing was applied using a 5-pixel sliding window approach. The myc-HspB3 signal in Figure 3F was enhanced in Fiji (Process > Math > Set Gamma to 0.5).

### Determination of minimal concentration threshold for condensate formation

To determine the average GFP-HspB2 signal intensity in cells that either did or did not contain condensates, mid-Z-stack optical sections were selected and duplicated in Fiji (Image > Duplicate). This duplicate was converted to 8-bit before a threshold was applied (Image > Adjust > Auto Threshold > Select ‘Moments’ with default setting). The thresholded, binary duplicate was normalized to 65535 (16 bit) and subtracted from the original image using Process > Image Calculator, and the mean intensity value for the GFP-HspB2 signal was determined for individual cells, correcting for the area of the cell that was removed in the previous step. Throughout sample processing, imaging, and image manipulations, the same settings were used for all samples, thus enabling direct comparison between cells.

### Determination of circularity of condensates and aggregates

To assess circularity of HspB2 condensates and aggregates, mid Z-stack optical sections were selected and converted to 8-bit. Bernsen local thresholding was performed using Fiji (Image > Adjust > Auto Local Threshold > Bernsen [radius = 100; parameter 1 = 100; parameter 2 = default]). Subsequently, area and circularity of thresholded structures was determined in Fiji (Analyze > Analyze particles [default settings, except size (µm^2^) set to 0.08 – infinity).

### Automated analysis of HspB2 condensate and aggregate size and localization

For the computational image analysis, python (>3.7, Delaware USA) was used. For the detection of the cells as well as the aggregates and condensates, images were thresholded with a global Otsu filter (76), after which a cut-off parameter was introduced. The obtained threshold was modified by multiplying its value with the cut-off value to remove weaker signals around the periphery of the aggregates and condensates. Finally, the remaining spurious noise in the image was removed (isolated pixel islands, the filter removes pixels when five neighbouring pixels are zero, the pixel of interest is set to zero as well). These steps generated two parameters (a cut-off and noise removal), which determined the size and shape of the detected aggregates and cells. To estimate the size of the aggregates defined in number of pixels, and to label individual aggregates, all possible graphs of distance for the nodes (all nonzero pixels) in the image matrix were created (77). Subsequently overlap with the DAPI signal was determined to assess whether the aggregates are located within the nucleus. To justify the final choice for the image analysis parameters (cut-off = 2.25 and noise filter = 5) the filtered images and data analysis for a range of relevant parameter values are shown in Figure S4. The code can be found on the https://github.com/huckgroup.

### Flow cytometry proliferation assay

For each condition, 1 x 10^5^ HEK293 cells were seeded in a single well of a 12-wells plate. After overnight incubation, cells were incubated in 100 nM CellTrace violet (Invitrogen, C34557) in PBS for 20 minutes at 37 °C. Subsequently, cells were washed twice with culture medium and fresh culture medium was added. Where specified, cells were transfected with indicated transfection mixes directly after incubation with the CellTrace violet solution. As a control, an empty pIRES vector not encoding HspB2 or HspB3 was used. At indicated time points, cells were washed with PBS and fixed in 4% PFA for 20 minutes. Afterwards, cells were washed once with PBS, resuspended in 200 µL FACS buffer (0.1% BSA, 0.05% NaN_3_, 0.5 mM EDTA in PBS), and stored at 4 °C awaiting FACS analysis. Flow cytometry was performed on a FACSVerse (BD Biosciences) flow cytometer. CellTrace violet was detected using the 405nm laser line and the 448/45 filter, GFP-HspB2 was detected using the 488nm laser line and the 510/20 filter. For the plots of HspB2-transfected cells shown in Figure S2B, only GFP-positive cells were analyzed to correct for variations in transfection efficiency.

### Fractionation of soluble and insoluble fractions

Cells grown in 12 wells plates were harvested using trypsin and pelleted by centrifuging at 1500 x *g* for 5 minutes at room temperature. Subsequently, cells were washed once with PBS, split into two replicate samples, and pelleted as before. The cell pellet of one replicate was lysed directly in 9 µL 4x SDS sample buffer (62.5 mM Tris-HCl [pH 6.8], 2% SDS, 5% β-mercaptoethanol, 10% glycerol, 0.005% bromophenol blue) to serve as total fraction. The other replicate was lysed in 25 µL RIPA lysis buffer (50 mM Tris-HCl [pH 7.5], 150 mM NaCl, 0.1% SDS, 0.5% sodium deoxycholate, 1% Triton X-100, 1x cOmplete protease inhibitor cocktail [Roche-11697498001], 1 mM PMSF), incubated on ice for 10 minutes and centrifuged for 10 minutes at 21100 x *g* at 4 °C. Subsequently, the supernatant was combined with 9 µL 4x SDS sample buffer to constitute the soluble fraction, while the insoluble pellet was washed once in RIPA buffer before adding 25 µL milliQ and 9 µL 4x SDS sample buffer. All samples were subsequently heated for 10 minutes at 95 °C.

### Western blotting

Samples were size separated on 12.5% polyacrylamide gels, transferred into nitrocellulose membranes and blocked with 5% w/v nonfat dry milk in PBS for 30 minutes at room temperature. In cases where membranes were stained for biotin, 3% BSA in TBST was used for blocking and as diluent for subsequent antibody incubations. After blocking, membranes were incubated for 1 hour at room temperature with indicated primary antibodies (rabbit-anti-HspB2 [generated in our laboratory; 1:1000], rabbit-anti-HspB3 [generated in our laboratory; 1:1000] (73), mouse-anti-tubulin-β (DSHB cat# E7, RRID:AB_528499; 1:1000]) diluted in 2.5% w/v nonfat dry milk in PBST. After extensive washing using PBST, membranes were incubated for 1 hour with secondary antibodies (goat-anti-rabbit-IRDye680 [Li-Cor Biosciences cat# 926-68071, RRID:AB_10956166], goat-anti-mouse-IRDye800 [Li-Cor Biosciences cat# 926-32210, RRID:AB_621842]) both at 1:10000 in 2.5% w/v nonfat dry milk in PBST. Afterwards, membranes were washed extensively in PBST again and imaged using the Li-Cor Odyssey CLx. For quantification of western blot signals in Figure 2H, intensity values were normalized using the tubulin-β signals of total samples to account for minor variations during cell culture, lysis and gel loading. Subsequently, the amount of HspB2 in the insoluble pellet fraction was calculated and expressed as a percentage relative to the HspB2 signal in the total sample.

### APEX2 proximity labelling

Proximity labelling was performed as described in (22). In brief, the APEX2-tag was amplified from the APEX2-NLS plasmid gifted by Alice Ting (Addgene plasmid # 124617) and introduced into pcDNA5 expression vectors, containing genes encoding HspB2, HspB5-WT, -STD/RG, and nuclear export/nuclear localization signals as indicated. Subsequently, these plasmids were transfected into HEK293 cells using PEI as described previously. 24 hours after transfection, cells were incubated in 500 µM biotin-phenol (BP) in full DMEM for 45 minutes at 37 °C. Next, a fresh 100x stock of 100 mM H_2_O_2_ in PBS was prepared, and H_2_O_2_ was directly added to the BP-solution to a final concentration of 1 mM. Cells were incubated for 1 minute, after which the labelling solution was removed. Cells were washed three times in quencher solution (10 mM sodium ascorbate, 5 mM trolox and 10 mM sodium azide in PBS) before fixation or cell lysis.

### Preparation of lysates for mass spectrometry

Plasmids encoding indicated APEX-tagged transgenes and untagged genes were transfected into HEK293 cells. For each lysate, 70 µg total plasmid DNA was used to transfect two T175 flasks seeded with HEK293 cells at approximately 70% confluency (See Figure S4A for exact composition of transfection mixes). 24 hours after transfection, APEX-proximity labelling was performed as above. After labelling, medium was removed and cells were washed twice with quencher solution, and twice with PBS, followed by a final wash with quencher solution. Each wash was performed for 1 minute, using 25 mL of wash solution. After the final wash, cells were harvested in quencher solution (10 mL per T175) using a cell scraper, transferred to 50 mL Falcon tubes and placed on ice. After all samples were harvested, cells were centrifuged at 3000 x *g* for 20 minutes at 4 °C. Cell pellets were resuspended in 1 mL PBS, transferred to Eppendorf tubes, and centrifuged for 2 minutes at 1500 x *g*. Cell pellets were resuspended in 120µL lysis buffer 1 (1% SDS, 1 mM DTT, 1x complete protease inhibitors, 1mM PMSF, 10 mM sodium ascorbate, 5 mM trolox, 10 mM sodium azide) and heated at 90 °C to promote solubilization of aggregated proteins. Subsequently, lysates were sonicated three times using a 30 sec on/30 sec off protocol. Next, lysates were diluted by adding 1080 µL lysis buffer 2 (56 mM Tris-HCl [pH 7.5], 167 mM NaCl, 0.55% NP-40, 0.55% sodium deoxycholate, 1x complete protease inhibitors, 1mM PMSF, 10 mM sodium ascorbate, 5 mM trolox, 10 mM sodium azide) and sonicated as before. Lysates were cleared by centrifugation at 21100 x *g*, 4 °C for 10 minutes. Supernatants were taken and dialyzed using Slide-A-Lyzer (Thermo, #66332) cassettes to remove free biotin-phenol. Briefly, the membranes were hydrated, after which the samples were loaded and dialyzed in 3 L dialysis buffer (50 mM Tris-HCl [pH 7.5], 150 mM NaCl, 1 mM DTT) for 2 hours at 4 °C. After 2 hours, dialysis buffer was refreshed and samples were dialyzed for an additional 2 hours. Subsequently, dialysis buffer was refreshed once more, and samples were incubated overnight. The following day, lysates were recovered, transferred to Eppendorf tubes and glycerol was added to a final concentration of 10%. After setting aside small fractions of each sample for Bradford assay and a small scale streptavidin enrichment test, lysates were snap-frozen in liquid nitrogen and stored at −80 °C. Protein concentration was determined using Bradford assay.

### Streptavidin enrichment and on-bead digestion

Throughout streptavidin enrichment and washing procedures, samples were kept on ice as much as possible. Streptavidin Sepharose High Performance beads (Cytiva, #17-5113-01) were equilibrated in IP-buffer (50 mM Tris-HCl [pH 7.5], 150 mM NaCl, 0.5% NP-40, 0.5% sodium deoxycholate, 0.1% SDS, 0.5 mM DTT, 1 x cOmplete protease inhibitor cocktail, 1 mM PMSF, 10 mM sodium ascorbate, 5 mM trolox, 10 mM sodium azide) three times, with centrifugation at room temperature for 2 minutes at 2000 x *g* in between. For each streptavidin enrichment, 15 µL bead slurry was used. Purification was performed in triplicate by combining 400 µg protein lysate in IP-buffer and equilibrated bead suspension in a total volume of 500 µL per replicate. In addition, 20µg ethidium bromide was added to prevent indirect, DNA-mediated interactions. Samples were incubated for 90 minutes at 4 °C with end-over-end rotation. Subsequently, beads were washed twice with wash buffer (25 mM Tris-HCl [pH 7.5], 150 mM NaCl, 1% NP-40, 1% sodium deoxycholate, 0.1% SDS, 0.5 mM DTT, 1 x cOmplete protease inhibitor cocktail, 1mM PMSF), twice with 1% NP-40 in PBS (MS-grade) and twice with PBS (MS-grade). From this point onwards, all incubations and centrifugation steps were performed at room temperature. The final wash was removed completely using an insulin syringe (BD cat# 324825, gauge 30), beads resuspended in 50 µL freshly prepared elution buffer 1 (2 M urea, 100 mM Tris-HCl [pH 8.0], 10 mM DTT) and incubated for 20 minutes in a thermoshaker set to 1250 rpm. Iodoacetamide was added to a final concentration of 50 mM and samples were incubated for an additional 10 minutes on the thermoshaker. After addition of iodoacetamide, samples were kept in the dark as much as possible. Subsequently, 0.25 µg trypsin (Promega, V5111C) was added and bead suspensions were incubated for approximately 150 minutes on the thermoshaker at 1250 rpm. After centrifugation for 2 minutes at 2000 x *g*, the supernatant containing tryptic peptides was taken and transferred to a new Eppendorf tube. Next, 50 µL freshly prepared elution buffer 2 (2 M urea, 100 mM Tris-HCl [pH 8.0]) was added to the beads, which were subsequently incubated for 5 minutes at 1250 rpm on the thermoshaker. Again, beads were pelleted by centrifugation (2 minutes, 2000 x *g*), supernatant was collected and combined with the supernatant taken previously. Combined supernatants were incubated overnight at room temperature after addition of 0.1 µg trypsin. Samples were acidified by adding 10 µL trifluoroacetic acid (10%), purified and desalted on C18 StageTips (78) and stored at 4 °C until measurement on the mass spectrometer.

### Mass spectrometry

Peptides were separated through reverse phase nano-HPLC on an Easy-nLC 1000 (Thermo Fisher) coupled on-line to a Thermo Fisher Orbitrap Exploris 480 mass spectrometer. A 60 minute gradient of buffer B (80% acetonitrile, 0.1% formic acid) was applied and the mass spectrometer was run in Top20 mode. After fragmentation, peptides were added to a dynamic exclusion list for 45 seconds.

### Mass Spectrometry data analyses

The RAW data have been analyzed using MaxQuant version 1.6.6.0 (79) LFQ and iBAQ settings enabled, and deamidation (NQ) added as variable modification. The database was downloaded from UniProtKB in June 2017 and APEX protein sequences were added. The raw mass spectrometry data and Maxquant output have been deposited to the ProteomeXchange Consortium via the PRIDE (80) partner repository with the dataset identifier PXD035040. Raw MS dataset identifiers of individual triplicates are indicated in Figure S5A.

Mapped mass spectrometry data was processed using Perseus 1.5.0.15 (81). In brief, identified proteins were filtered for contaminants and reverse hits, after which LFQ-values were log2-transformed. Samples were grouped into triplicates and filtered to contain three valid values in at least one group of replicates. Missing values were subsequently imputed assuming normal distribution to allow statistical analyses. Volcano plots were generated in Perseus and replotted in GraphPad Prism 8. GO-term analysis of proteins enriched in HspB2 condensates and HspB5-STD/RG aggregates was performed on proteins in the 95^th^ percentile with *P* < 0.01 in the appropriate comparison. Similarly, for the analysis of proteins depleted in HspB2 condensates, the 5% most depleted proteins with a *P*-value < 0.01 were selected. To investigate GO-term enrichment of factors enriched in HspB2:HspB3 aggregates, proteins in the 90^th^ percentile relative to both APEX-NES and -NLS controls were selected. These datasets were then overlapped and only those proteins present in both comparisons were selected for GO-term enrichment analysis. GO-term enrichment analysis (82–84) was performed by testing lists of selected proteins against the human proteome reference list on http://geneontology.org/. Only enriched terms with an FDR *P* < 0.05 are shown.

## SUPPLEMENTAL INFORMATION for

**Figure S1.**
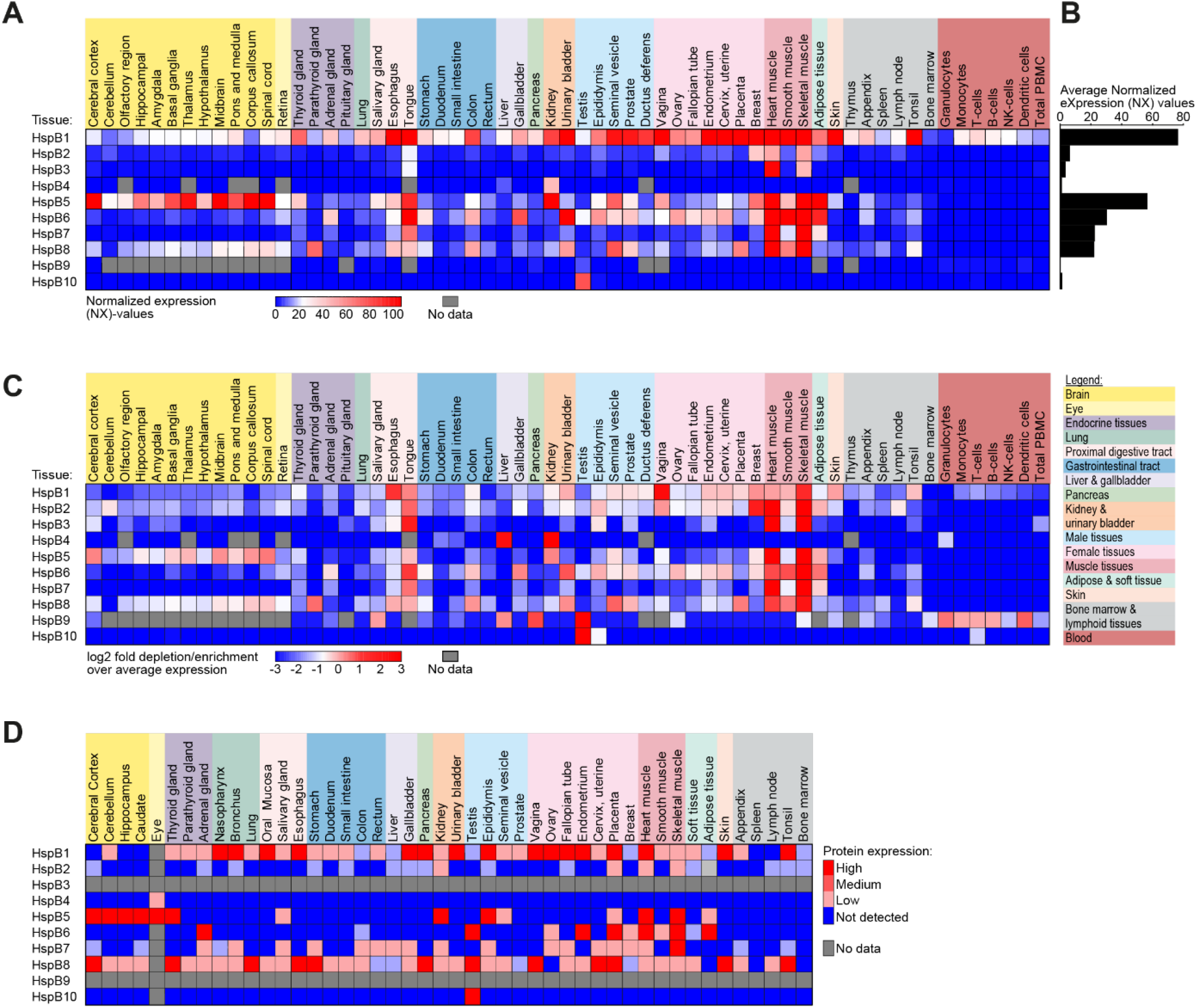
Tissue-specific expression pattern of human small heat shock proteins. **(A)** Heat-map depicting normalized expression (NX) values of all ten human small heat shock proteins (HspB1-10) in indicated tissues. Grey boxes are used to indicate sHsp-tissue combinations for which no data was available. **(B)** Average normalized expression values for individual members of the small heat shock protein family across all tissues indicated in (A). **(C)** Heat-map indicating the log2 fold enrichment (red) and -depletion (blue) of sHSP-expression in individual tissues over the average expression level plotted in (B). **(D)** Protein expression of HspB1-10 in indicated tissues, as deducted from immunohistochemical staining data. Expression was scored as high, medium, low, or not-detected. All data used to generate the figures in (A-D) was extracted from www.proteinatlas.org.

**Figure S2.**
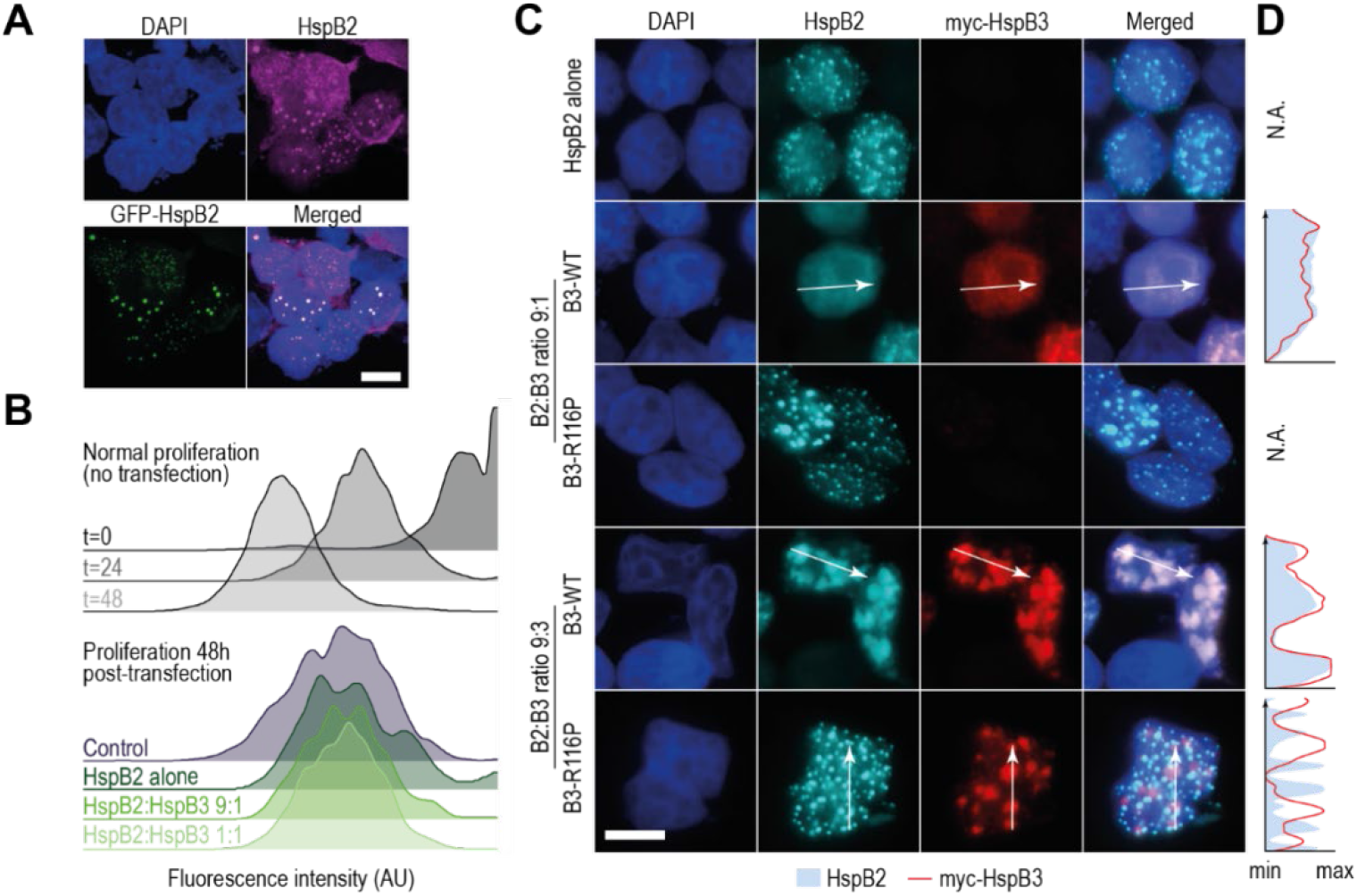
HspB3 affects subcellular localization of HspB2, but does not impact cell proliferation. **(A)** Confocal imaging of HEK293 cells transfected with GFP-tagged and untagged HspB2 at a 1:8 ratio, co-stained with an antibody targeting HspB2. The scalebar denotes 10µm (also for (C)). **(B)** Flow cytometry proliferation assay of untransfected cells at indicated time-points (top) and cells transfected with indicated plasmid mixes 48 hours post-transfection (bottom). Transfected cells proliferate slower compared to untransfected cells, regardless of what ratio of plasmids was used. Importantly, cells transfected with an empty control plasmid that does not encode HspB2 or HspB3 (purple) proliferate at a similar rate as cells transfected with HspB2 and HspB3 encoding plasmids, indicating the observed reduction in proliferation speed is a general transfection effect, and does not result from condensation or aggregation of HspB2. **(C)** Confocal images of cells expressing HspB2 and myc-tagged wildtype (WT) and mutant (R116P) HspB3. The scalebar denotes 10µm and the arrows indicate the direction along which the histogram in (D) was determined. **(D)** Histograms depicting HspB2 and myc-HspB3 signal intensities along the lines indicated in (C).

**Figure S3.**
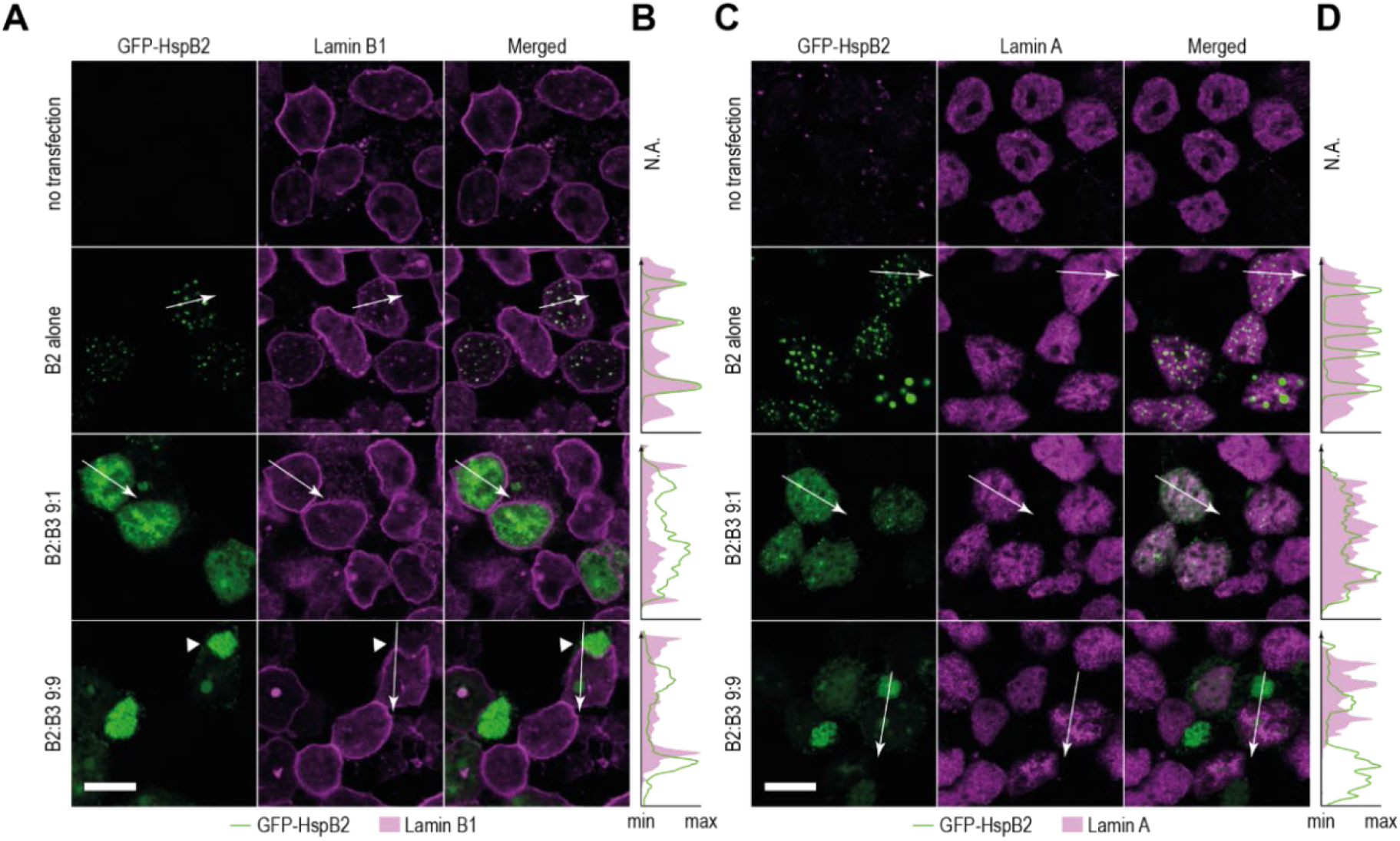
Lamin B1 colocalizes with HspB2 in nuclear condensates. **(A)** Confocal imaging of HEK293 cells transfected with HspB2 and HspB3 at indicated ratios, co-stained with an antibody against Lamin B1. The arrows indicate the line and direction along which the colocalization histograms in (B) were drawn. The solid arrowhead indicates a cytoplasmic aggregate that distorts the nuclear shape. The scalebar denotes 10μm. **(B)** Signal intensity histogram depicting the GFP-HspB2 and Lamin B1 signals along the lines shown in (A). **(C-D)** As (A-B), but with cells co-stained using an antibody targeting Lamin A.

**Figure S4.**
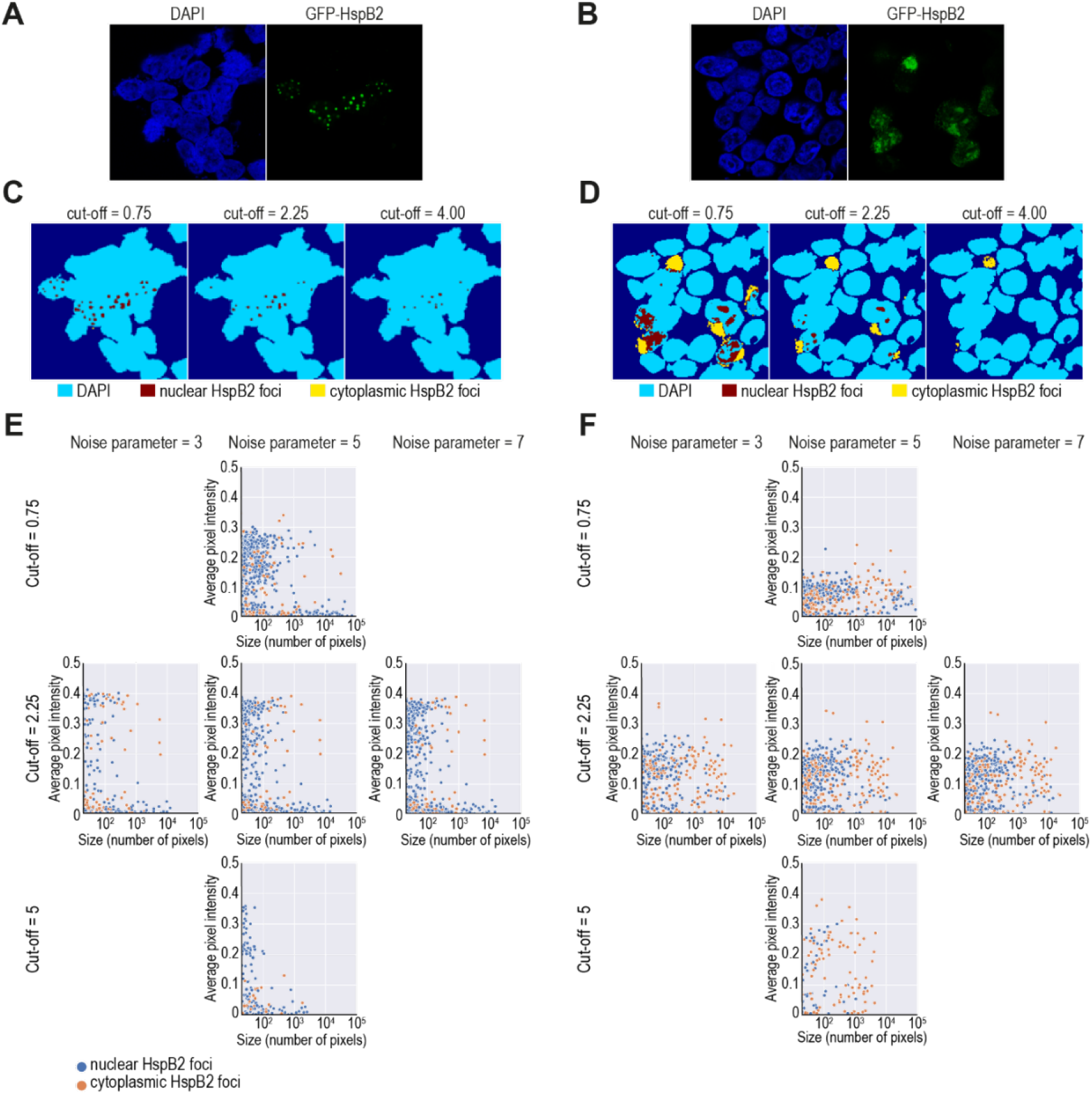
Justification of thresholds for signal quantification. **(A-B)** Raw DAPI and GFP-HspB2 signal in cells expressing HspB2 alone (A) or HspB2 and HspB3 at a 1:1 ratio (B), which underly the segmentation shown in (C) and (D), respectively. **(C-D)** Segmentation and HspB2 foci identification, using three different values for the threshold parameter in cells expressing HspB2 alone (C) or HspB2 and HspB3 at a 1:1 ratio (D). **(E-F)** Scatter plots depicting the size, average pixel intensities, and subcellular localization of HspB2 foci in cells expressing HspB2 alone (E) or HspB2 and HspB3 at a 1:1 ratio (F), using different values for threshold and noise parameters.

### SUPPLEMENTAL TEXT

To ensure that our automated image analysis script did not introduce a bias in the identification of HspB2 condensates and aggregates, we tested the effect of a range of cut-off and noise filter values. We took images from cells expressing HspB2 alone (Figure S4A) or HspB2 and HspB3 at a 1:1 ratio (Figure S4B), and filtered at a mid-stack Z-position using different cut-off values for the global thresholding method (ranging from 0 - 4). Here, we found that, at a value of 0.75, the filter uncovers a large number of nuclear HspB2 foci in cells expressing HspB2 alone (Figure S4C). The abundance of identified foci decreases as the cut-off value increases, and at values > 4, barely any HspB2 foci are detected after the thresholding step. In addition, we also assessed the effect of various thresholding values in cells transfected with HspB2 and HspB3 at equimolar ratios. As before, abundant HspB2 foci were detected at low cut-off values, and most foci were lost step at cut-off values exceeding 4 (Figure S4D).

We subsequently tested whether the threshold parameter influenced the analysis outcome, using our entire dataset. Altering the cut-off values changed the results quantitatively, yet, qualitatively the conclusions that can be drawn from this analysis do not change within the detection range of the threshold parameter. Briefly, for all cut-off values, the HspB2 foci in cells expressing HspB2 alone are relatively small and nuclear (Figure S4E), whereas foci in HspB2:HspB3 1:1 cells are larger and distributed across the nucleus and cytoplasm (Figure S4F). Similarly, altering the noise parameter did not change the result qualitatively to any significant degree (Figure S4E-F). Based on these observations, we opted for a global threshold cut-off value of 2.25 and a noise filter value of 5 for our final analysis. At www.gitlab.com/hucklab we provide a demo script, where the effect of the image analysis parameters can be tested using 2 example images.

**Figure S5.**
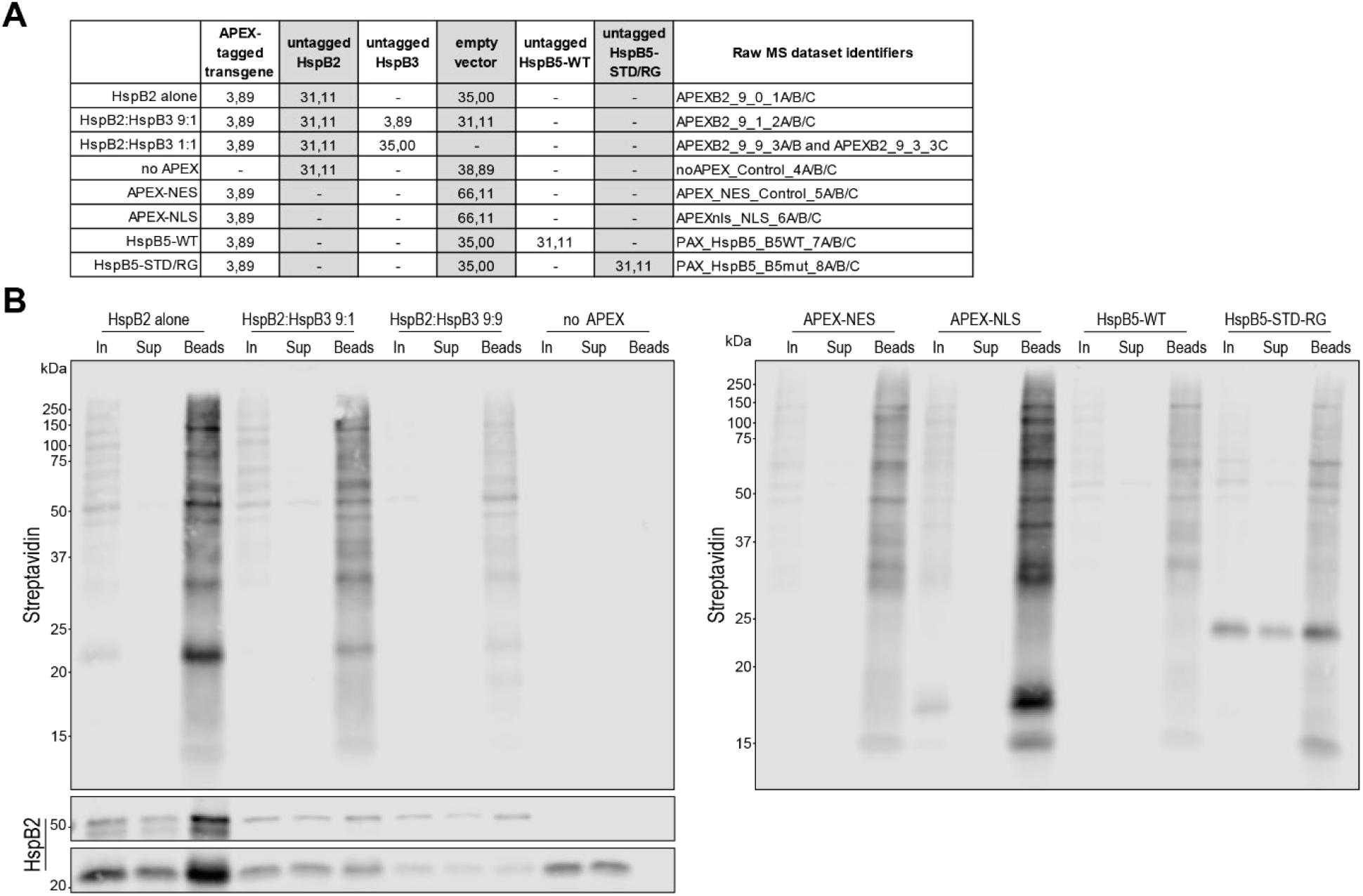
Transfection mixes and streptavidin enrichment for mass spectrometry experiments. **(A)** The amount of various plasmid-encoded transgenes (in µg) that was transfected for mass spectrometric analysis of the APEX-labelled interactome in indicated samples. **(B)** Western blot showing protein biotinylation using a fluorophore coupled Streptavidin in input (In) and supernatant (Sup) samples, as well as biotinylated proteins enriched on streptavidin-coated beads (Beads). Untagged and APEX-tagged HspB2 was detected using an antibody against HspB2.

**Figure S6.**
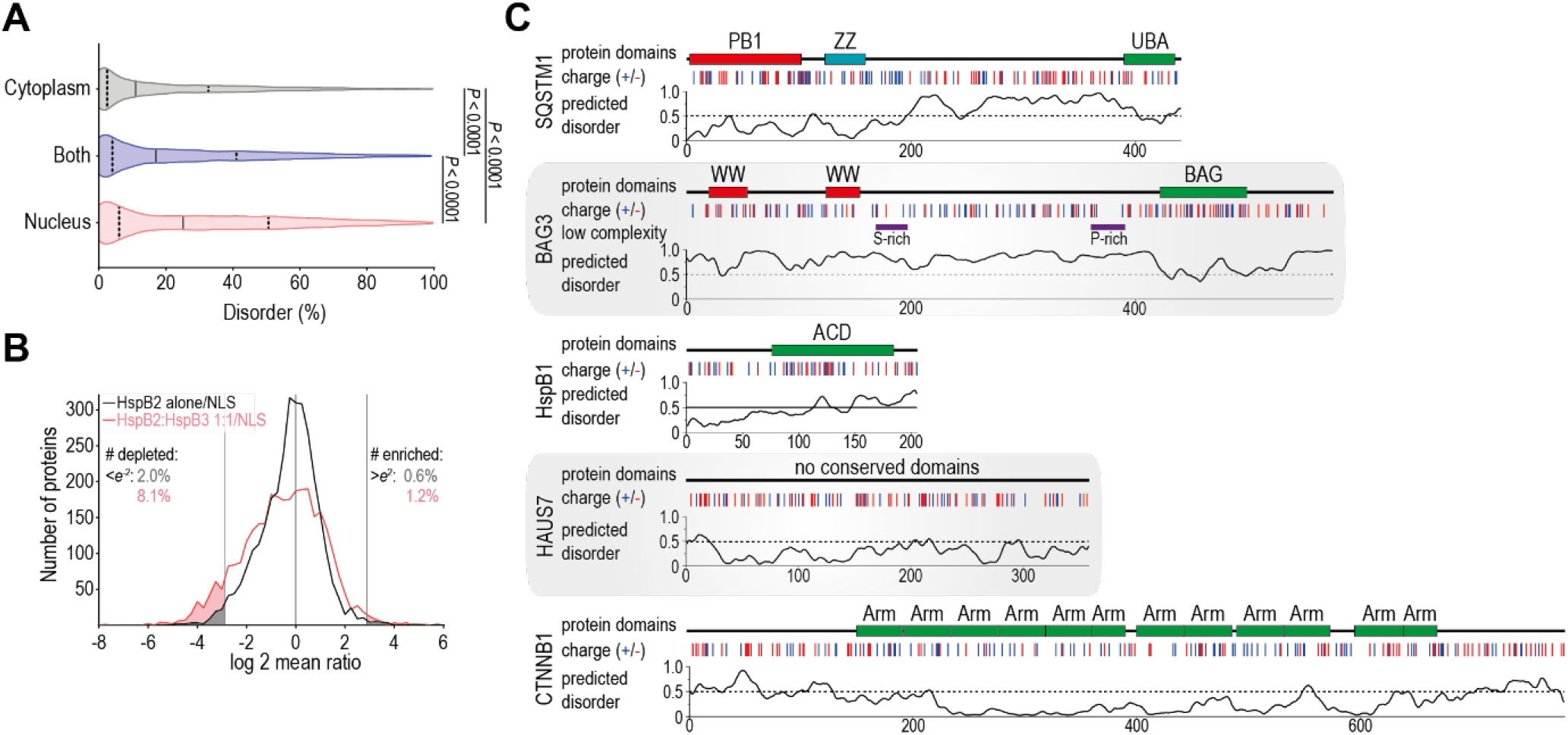
Proximity labelling uncovers that entrapment of interacting proteins in HspB2 foci is limited. **(A)** Violin plot depicting the distribution of the predicted percentage of disordered residues in proteins localized to the nucleus (red; n=5376), cytoplasm (grey, n=4567), or both (blue, n=2952). The median and quartiles are indicated by solid and dashed lines, respectively. Statistical significance was determined using a Kruskal-Wallis test followed by Dunn’s multiple comparisons test. **(B)** Histogram depicting the distribution of log2-fold changes of all identified proteins in HspB2 alone/NLS (black) and **(A) (A)** HspB2:HspB3/NLS (red) comparisons. Lines indicate *e*^−2^ and *e*^2^ cut-off values and the numbers indicate the fraction of proteins that are depleted or enriched beyond these cut-offs. **(C)** Schematic representation of the domain structure, charge distribution, low complexity regions, and predicted disorder in the five proteins used for validation. Protein domain organization was determined by combining information from the Uniprot website and predictions from the NCBI conserved domain database (www.ncbi.nlm.nih.gov/Structure/cdd/wrpsb.cgi), using default settings. Positions of positively charged (lysine, arginine, and histidine) and negatively charged (aspartic acids and glutamic acid) amino acid residues are indicated in blue and red, respectively. Low complexity regions were determined using PlaToLoCo (39), showing only those regions identified by the SEG-intermediate and SEG-strict methods (40). Protein disorder plots were generated with IUpred3 using default settings (41). PB1 - Phox and Bem1 domain; ZZ – ZZ-type zinc finger; UBA - ubiquitin-associated domain; WW – WW-domain; BAG – BAG-domain; ACD - α-crystallin domain; Arm - armadillo repeat.

**Figure S7.**
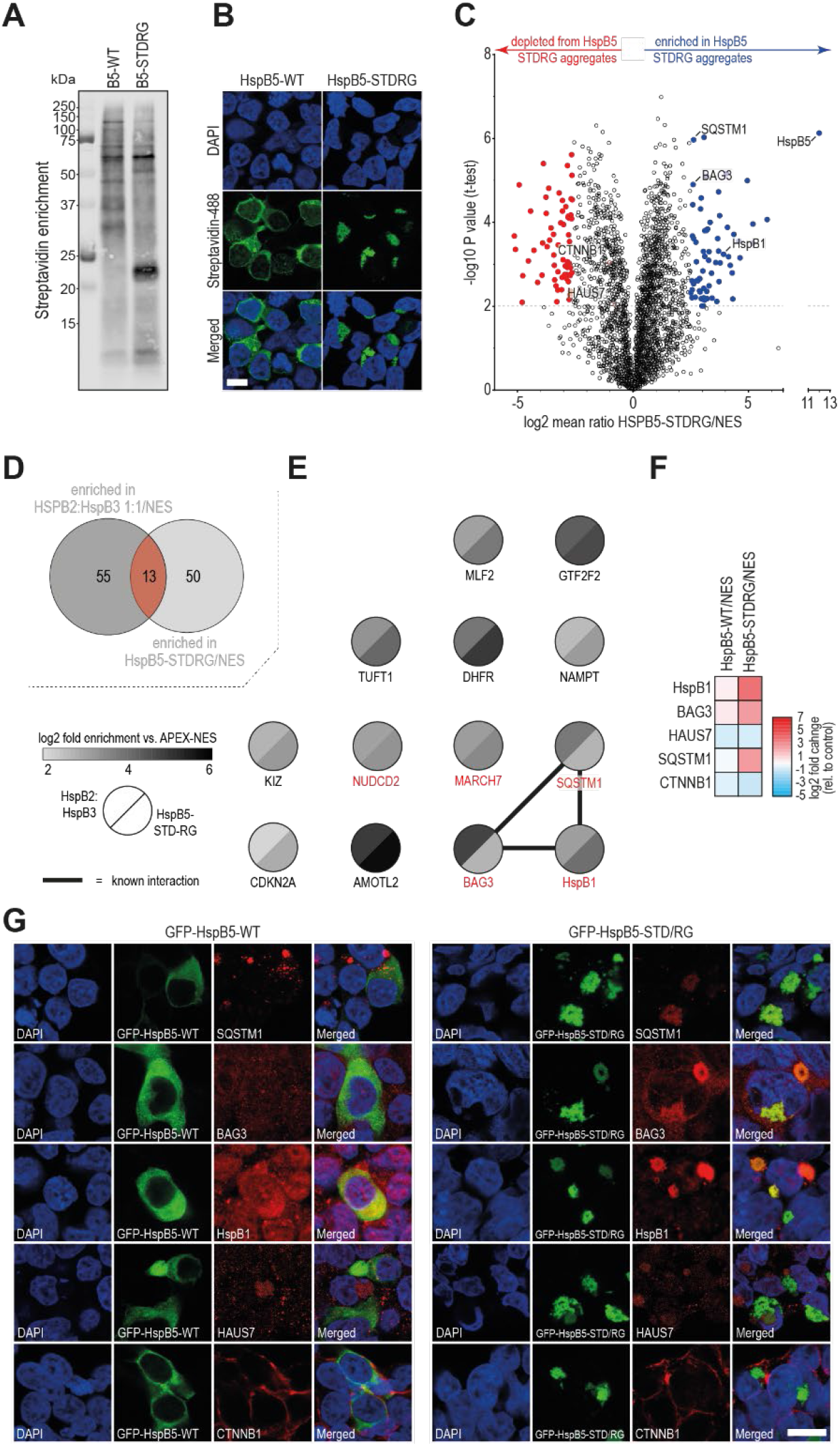
A similar set of autophagy factors is recruited to two types of sHSP aggregates. **(A)** Streptavidin-blotting of enriched biotinylated proteins from HEK293 cells expressing APEX-tagged wildtype (WT) and mutant (STD/RG) HspB5. **(B)** Confocal microscopy images of biotinylation in APEX-labelled HEK293 cells expressing wildtype (WT) and mutant (STD/RG) HspB5. Scale bar is 10µm. **(C)** Volcano plot showing proteins enriched (blue) and depleted (red) in HspB5-STD/RG proximity labelling relative to APEX-NES. The top 5% enriched and depleted proteins with a *P*-value <0.01 are highlighted. **(D)** The overlap between the top 5% enriched factors with a *P*-value <0.01 in HspB2:HspB3 1:1 and HspB5-STD/RG datasets was determined to uncover common factors enriched in both datasets. **(E)** Schematic overview of the enrichments over APEX-NES of the 13 common factors identified in (D). Grey shading indicates the enrichment in HspB2:HspB3 (top left) and HspB5-STD/RG (bottom right) datasets. Black lines indicate known physical interactions, and red font indicates proteins linked to GO-terms associated with protein (mis)folding, ubiquitination, ubiquitin binding and/or autophagy. **(F)** Summary of the log2 fold changes of five candidate proteins in HspB5-WT and -STD/RG proximity labelling datasets. **(G)** Confocal microscopy images of the five candidate proteins, along with GFP-tagged HspB5-WT and -STD/RG. Scalebar indicates 10µm.

### SUPPLEMENTAL VIDEO LEGENDS

Video S1-S3 and Table S1 can be accessed through the following link: https://surfdrive.surf.nl/files/index.php/s/caqnxXYLWrMTgKl

**Video S1. Live imaging of HEK293 cells transfected with a plasmid encoding HspB2**

HEK293 cells transfected with plasmids encoding GFP-tagged and untagged HspB2 (1:8 ratio) were visualized using the Leica DMi8 widefield microscope. Imaging stared ~16 hours after transfection and continued for approximately 15 minutes with an imaging frequency of 30 frames per minute. Dynamic movement of circular HspB2 condensates can be seen in multiple cells. The cell depicted in Figure 1C and Video S2 is located in the bottom-right corner.

**Video S2. Fusion of HspB2 condensates**

Live imaging showing fusion of two HspB2 condensates in the nucleus of a HEK293 cell transfected with plasmids encoding GFP-tagged and untagged HspB2 (1:8 ratio). The clock in the top-left corner indicates the time passed the start of imaging in mm:ss. The first image was taken approximately 16 hours after transfection.

**Video S3. The development of HspB2 condensates and HspB2:HspB3 aggregates**

**(A-C)** Time-lapse videos of the development of HspB2 signal in HEK293 cells transfected with HspB2 alone (A), and HspB2:HspB3 at ratios 9:1 (B) and 9:3 (C). Cells were imaged every 15 minutes between ~5-24 hours post-transfection.

**Table S1. Overview of enriched and depleted proteins in APEX-proximity labelling datasets**

**(A)** List of proteins enriched in HspB2 condensates (top 5% enriched proteins with a P value <0.01 in the HspB2 alone/NLS comparison [blue in Figure 5A]).

**(B)** List of proteins depleted from HspB2 condensates (bottom 5% enriched proteins with a P value <0.01 in the HspB2 alone/NLS comparison [red in Figure 5A]).

**(C)** List of proteins enriched in HspB2:HspB3 aggregates: top 10% enriched in HspB2:HspB3 1:1/NLS and HspB2:HspB3/NES [red overlap in Venn diagram in Figure 6C]).

**(D)** List of proteins enriched in HspB5-STDRG aggregates (top 5% enriched proteins with a P value <0.01 in the HspB5-STDRG/NES comparison [blue in Figure S7C]).

**(E)** List of proteins depleted from HspB5-STDRG aggregates (bottom 5% enriched proteins with a P value <0.01 in the HspB5-STDRG/NES comparison [red in Figure S7C]).

### SUPPLEMENTAL DATASET 1

**Supplemental dataset 1.**
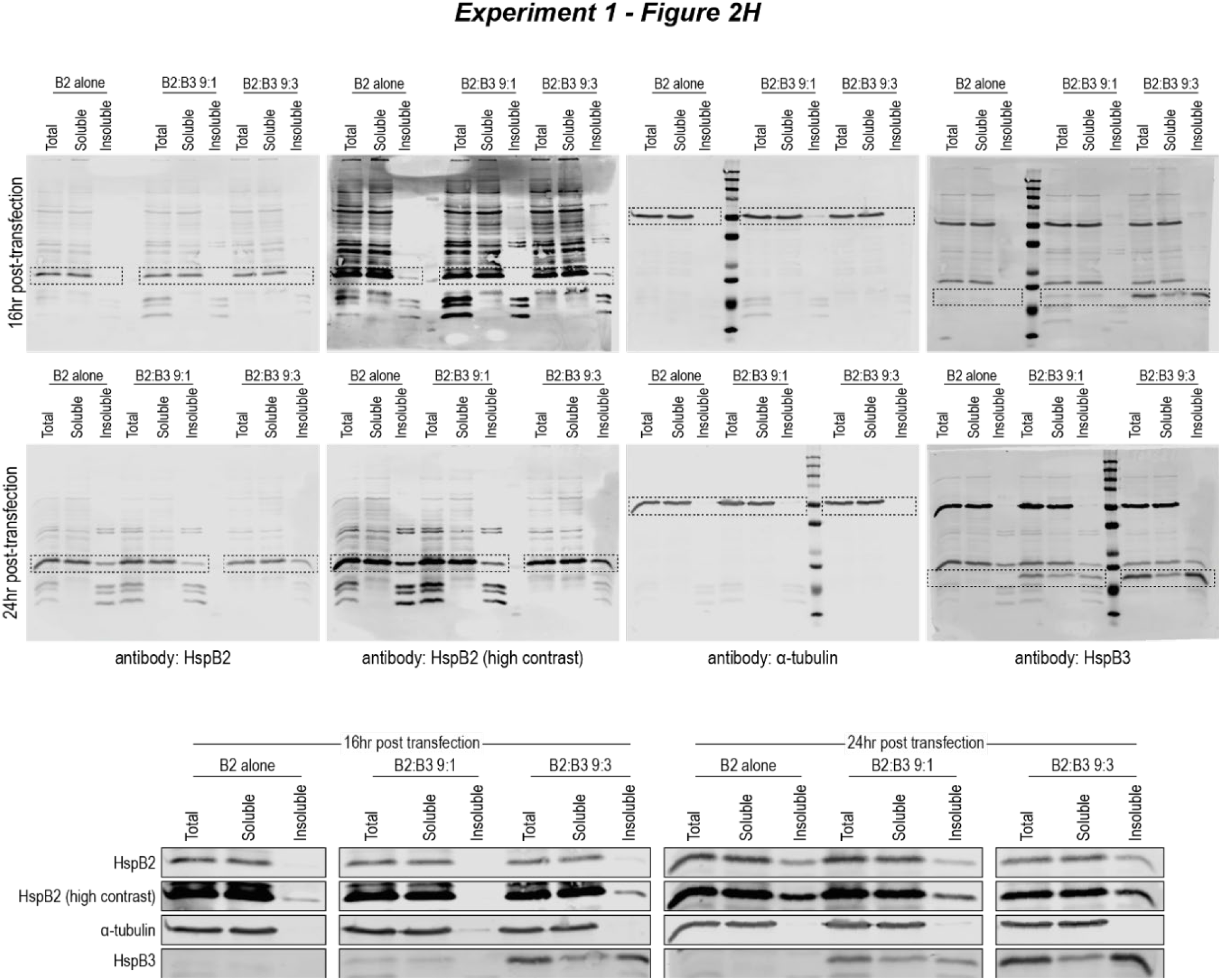
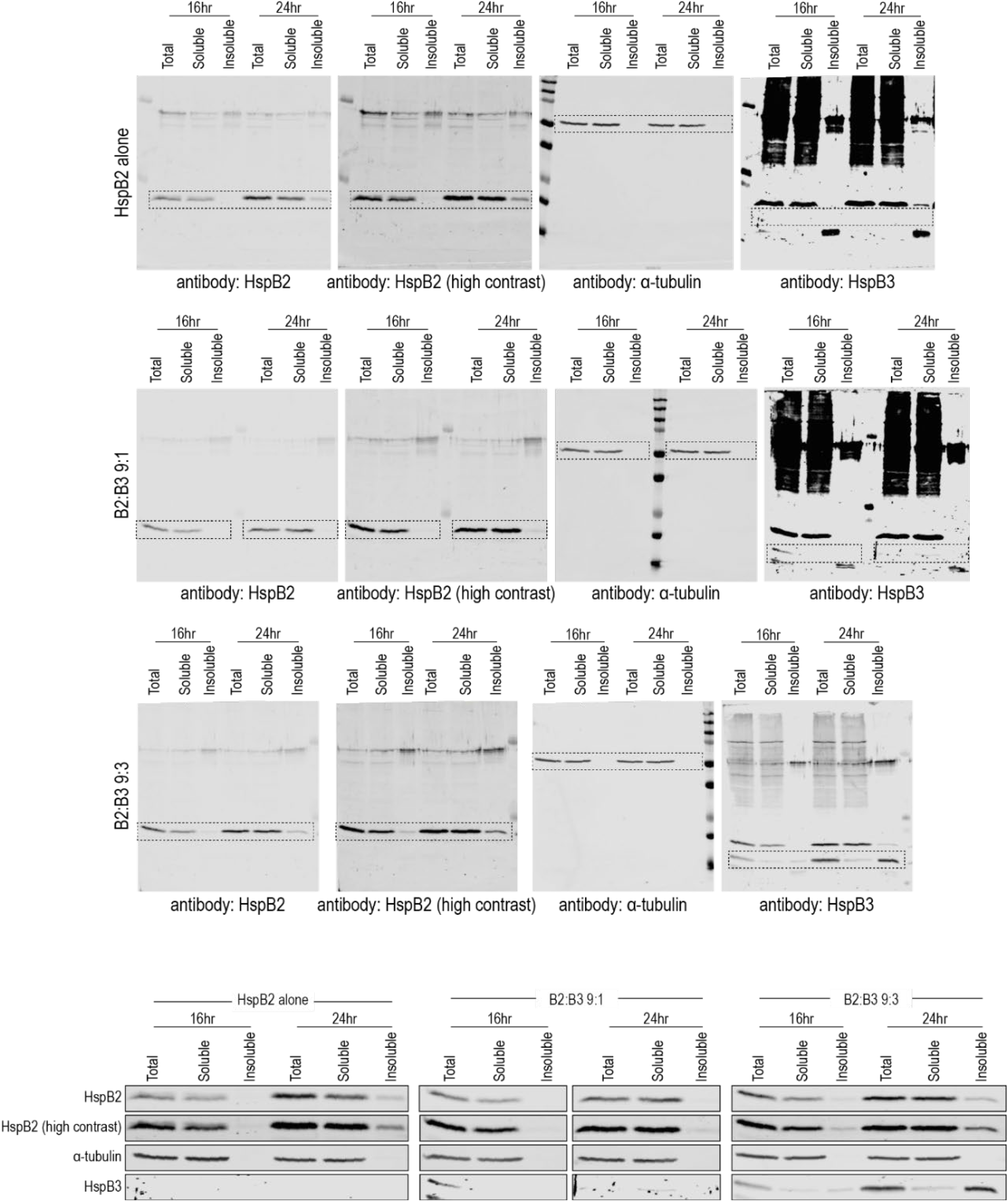
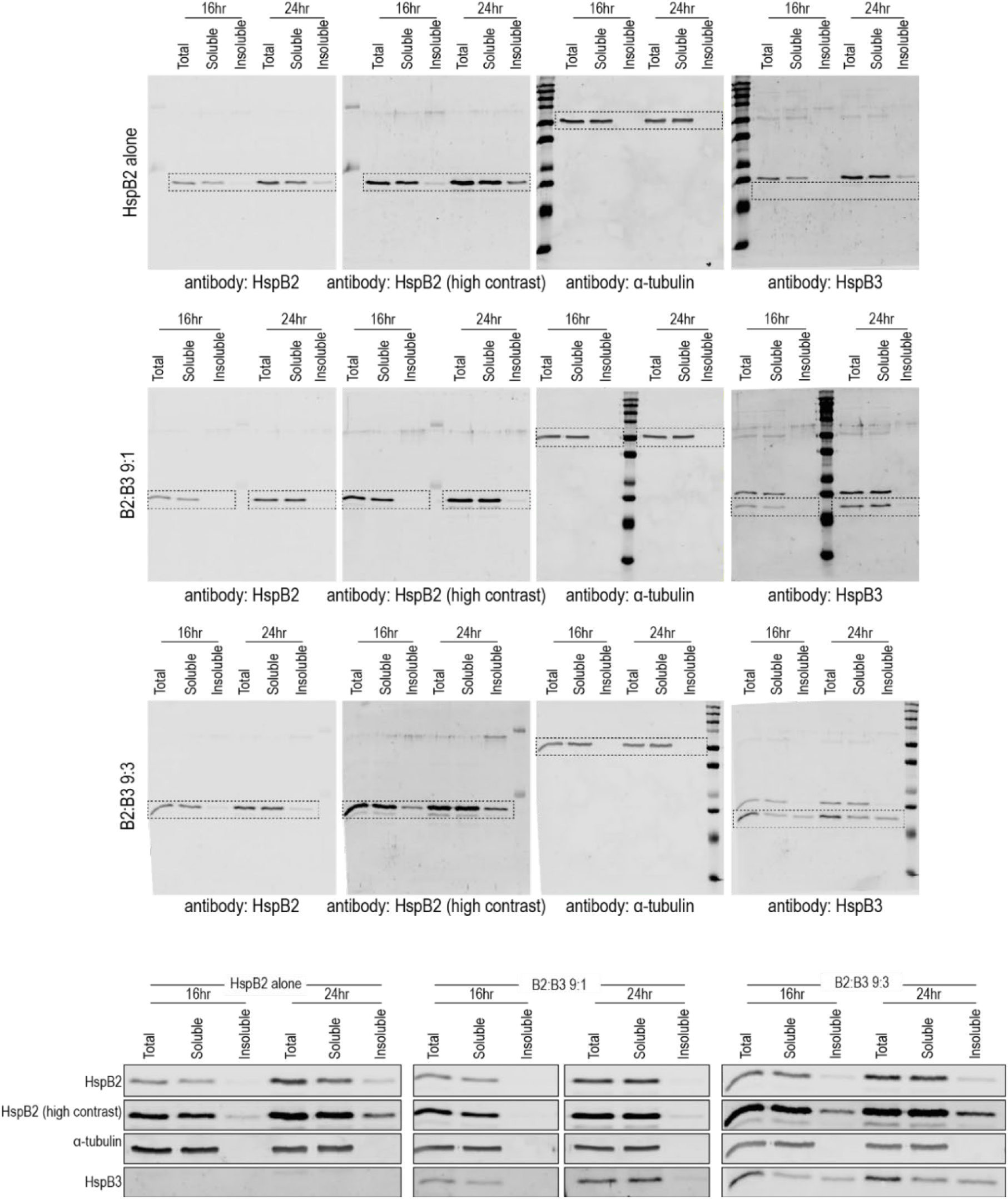
Western blot data underlying signal quantification shown in Figure 2H. HspB2, HspB3, and α-tubulin were detected in the soluble and insoluble fractions of lysates from HEK293 cells transfected with HspB2- and HspB3-expression vectors at indicated ratios. HspB2 signals were quantified and normalized using the α-tubulin signal in total lysates to correct for slight variations in cell numbers.

### SUPPLEMENTAL DATASET 2

**Supplemental dataset 2.**
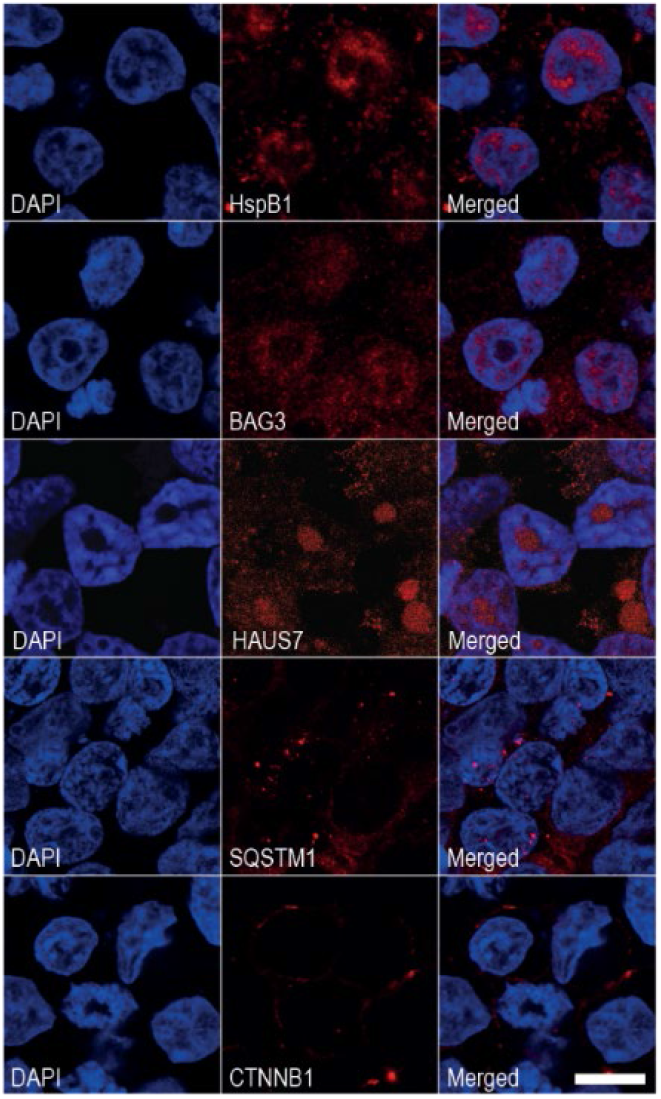
Baseline expression of HspB1, BAG3, HAUS7, SQSTM1, and CTNNB1. The five candidate genes used for validation of the proximity labelling dataset were detected to establish baseline subcellular distribution patterns in untransfected cells. Scalebar denotes 10µm.

## Notes

https://surfdrive.surf.nl/files/index.php/s/caqnxXYLWrMTgKl

### Competing Interest Statement

The authors have declared no competing interest.

